# ABCG transporters are involved in the accumulation of specialized metabolites in the oil bodies of *Marchantia polymorpha*

**DOI:** 10.1101/2025.10.27.684951

**Authors:** Takehiko Kanazawa, Tomoko Kawaguchi, Yuta Moriwaki, Kan-ichi Iwasaki, Takashi Ueda, Kenji Matsui

## Abstract

Bioactive specialized metabolites (SMs) are synthesized and sequestered in specific cellular compartments or organelles as a self-defense strategy against their intrinsic toxicity. Liverwort-specific oil bodies accumulate large amounts of SMs and contribute to chemical defense; however, the molecular mechanisms underlying SM sequestration in oil bodies remain largely unknown. Therefore, in this study, we focused on MpABCG1 and MpABCG36, which are ATP-binding cassette (ABC) protein family members localized to the oil bodies of liverwort *Marchantia polymorpha*. Sesquiterpene (thujopsene, chamigrene, and himachalane) accumulation was reduced in the Mp*abcg1* and Mp*abcg36* loss-of-function mutants. Notably, levels of the bisbibenzyls, marchantins C and A, were predominantly reduced in Mp*abcg1*, but not in Mp*abcg36*. Although the Mp*abcg1* mutant formed a number of oil bodies labeled with mCitrine– MpSYP12B (oil body membrane marker) comparable to that of the wild-type, the number of oil bodies stained with BODIPY 493/503, which has an affinity for lipophilic SMs, was reduced. This finding suggests that Mp*ABCG1* and Mp*ABCG36* mutations affect SM accumulation in the oil body but have little impact on oil body formation. Overall, our results highlight the involvement of Mp*ABCG1* and Mp*ABCG36* in the accumulation of SMs and/or their precursors in liverwort oil bodies.

## Introduction

Plants produce and accumulate various specialized metabolites (SMs), such as sesquiterpenoids, flavonoids, and alkaloids. Highly bioactive compounds that defend against herbivores and pathogens often exert toxic or harmful effects on plants themselves. Therefore, SM biosynthesis and accumulation are strictly, and often spatiotemporally, regulated (Blomstedt et al. 2016; Tissier 2012; Widhalm et al. 2015). SMs and/or their precursors synthesized in plastids or the cytosol are transported and stored in specific organelles and compartments, such as the vacuole and extracellular space, via membrane transporters and/or membrane trafficking systems (Ichino and Yazaki 2022).

Among bryophytes (hornworts, mosses, and liverworts), >90% of liverwort species possess a unique organelle called an oil body, in which various SMs accumulate (Asakawa et al. 2013; He et al. 2013; Romani et al. 2022; Suire et al. 2000; Tanaka et al. 2016). Oil bodies of liverworts differ from those of seed plants (also known as lipid bodies or oleosomes); the former are formed via redirection of the secretory pathway, whereas the latter arise via budding from a subdomain of the endoplasmic reticulum (ER) (Huang 1992; Kanazawa et al. 2020). In *Marchantia polymorpha*, oil bodies are formed in idioblastic “oil body cells,” which accumulate unique sesquiterpenes (e.g., thujopsene, chamigrene, and himachalene) and bisbibenzyls (marchantins C and A), as revealed via single-cell extraction using microcapillaries, followed by direct mass spectrometry (MS) analysis (Tanaka et al. 2016). Several transcription factors, including MpTGA, MpC1HDZ, MpMYC, MpERF13, and MpMYB02, act as key regulators of oil body cell differentiation, oil body formation and maturation, and SM-related enzyme expression (Gutsche et al. 2024; Kanazawa et al. 2020; Kubo et al. 2024; Peñuelas et al. 2019; Romani et al. 2020; Wang et al. 2023). However, the molecular mechanisms underlying SM accumulation in oil bodies remain unknown.

Existing knowledge on angiosperms suggests that sesquiterpenoids in liverwort oil bodies are biosynthesized via the cytosolic mevalonate (MVA) pathway from precursor isopentenyl diphosphate (IPP) through farnesyl diphosphate (FPP). In contrast, monoterpenoids are possibly synthesized via the plastid-localized 2-C-methyl-D-erythritol 4-phosphate (MEP) pathway (Vranová et al. 2013). Intermediates and/or final products are presumably transported into the oil body lumen across the phospholipid bilayer membrane. Marchantins are potentially biosynthesized from phenylalanine via the phenylpropanoid pathway (Friederich et al. 1999a). The condensation of dihydro-*p*-coumaroyl-CoA with three molecules of malonyl-CoA produces prelunularic acid, which is subsequently converted into bibenzyl lunularic acid. A putative cytochrome P450 enzyme catalyzes the oxidative coupling of two lunularic acid molecules to form marchantin C in an NADPH-dependent manner. A portion of marchantin C is further hydroxylated, possibly by another cytochrome P450 enzyme, to yield marchantin A (Friederich et al. 1999b).

As most cytochrome P450 enzymes are localized to the ER membrane and require reducing equivalents supplied by NADPH-dependent cytochrome P450 reductases (Achnine et al. 2004), marchantins are possibly biosynthesized in the cytosol (Friederich et al. 1999b). Their intermediates and/or final products are presumably transported to the oil body lumen across the oil body membrane. As molecular transport across biological phospholipid bilayers is generally mediated by membrane transporter proteins (Ichino and Yazaki 2022; Widhalm et al. 2015), one potential candidate for mediating SM transport to the oil body lumen is MpABCG1, which specifically localizes to the oil body membrane (Kanazawa et al. 2020).

ATP-binding cassette (ABC) transporters, present in all kingdoms, mediate the transport of various molecules, including metals, hormones, lipids, and phenolics. ABC transporters consist of either a single nucleotide-binding domain (NBD) fused with a single set of transmembrane domains (TMDs), referred to as “half” transporters, or two NBDs and two sets of TMDs, referred to as “full” transporters. ATP hydrolysis in NBDs provides the driving force for substrate translocation across membranes, whereas TMDs form a tunnel through which the substrates pass (Wilkens 2015). Phylogenetic analyses have revealed that ABC transporters in the plant lineage are classified into eight subfamilies: A–G and I (Verrier et al. 2008). Specifically, ABCG subfamily is markedly expanded in plants. Half ABCG transporters in plants are sometimes referred to as “white–brown complexes” because of their phylogenetic relationship with those involved in eye pigmentation in fruit flies (Gräfe and Schmitt 2021). For example, volatile terpenoids from petunia flowers are transported from the interior of petal cells to the atmosphere by a half ABCG transporter (Adebesin et al. 2017). Plants also possess a large group of full ABCG transporters that are absent in animals (Verrier et al. 2008). They form a distinct clade corresponding to the pleiotropic drug resistance subfamily and play various roles, including pathogen defense, diffusion barrier formation, and phytohormone transport, in plants (Gräfe and Schmitt 2021). Full ABCG transporters also participate in the deposition of cutin molecules and secretion of antifungal terpenes and alkaloids (Ichino and Yazaki 2022). Additionally, both half and full ABCG transporters are involved in monoterpene emission from the flowers of scented orchid species (Chang et al. 2023).

In this study, we focused on the oil body-localized transporters, MpABCG1 and MpABCG36. These two ABC transporters cooperatively contributed to the biosynthesis and accumulation of sesquiterpenes, with MpABCG1 playing predominant roles in the biosynthesis and accumulation of marchantins. Our findings provide new insights into the mechanisms underlying SM accumulation in the oil bodies of *M. polymorpha*.

## Results

### MpABCG1 and MpABCG36 are specifically expressed in oil body cells and localized to the oil body membrane in *M. polymorpha*

To investigate the mechanisms underlying the accumulation of sesquiterpenoids and marchantins in the oil bodies of *M. polymorpha*, we focused on MpABCG1, which localizes to the oil body membrane (Kanazawa et al. 2020). MpABCG1 is composed of two NBDs and two sets of TMDs, arranged in the order NBD–TMD–NBD–TMD, thus representing a full ABCG protein (Figure 1). *M. polymorpha* genome encodes 13 full and 23 half ABCG proteins (NBD–TMD; Figure 1) (Bowman et al. 2017). To clarify the phylogenetic relationships among full ABCG proteins, we conducted maximum-likelihood phylogenetic analysis and found that MpABCG36 (Mp2g14210) was phylogenetically close to MpABCG1 (Figure 2).

**Figure 1.**
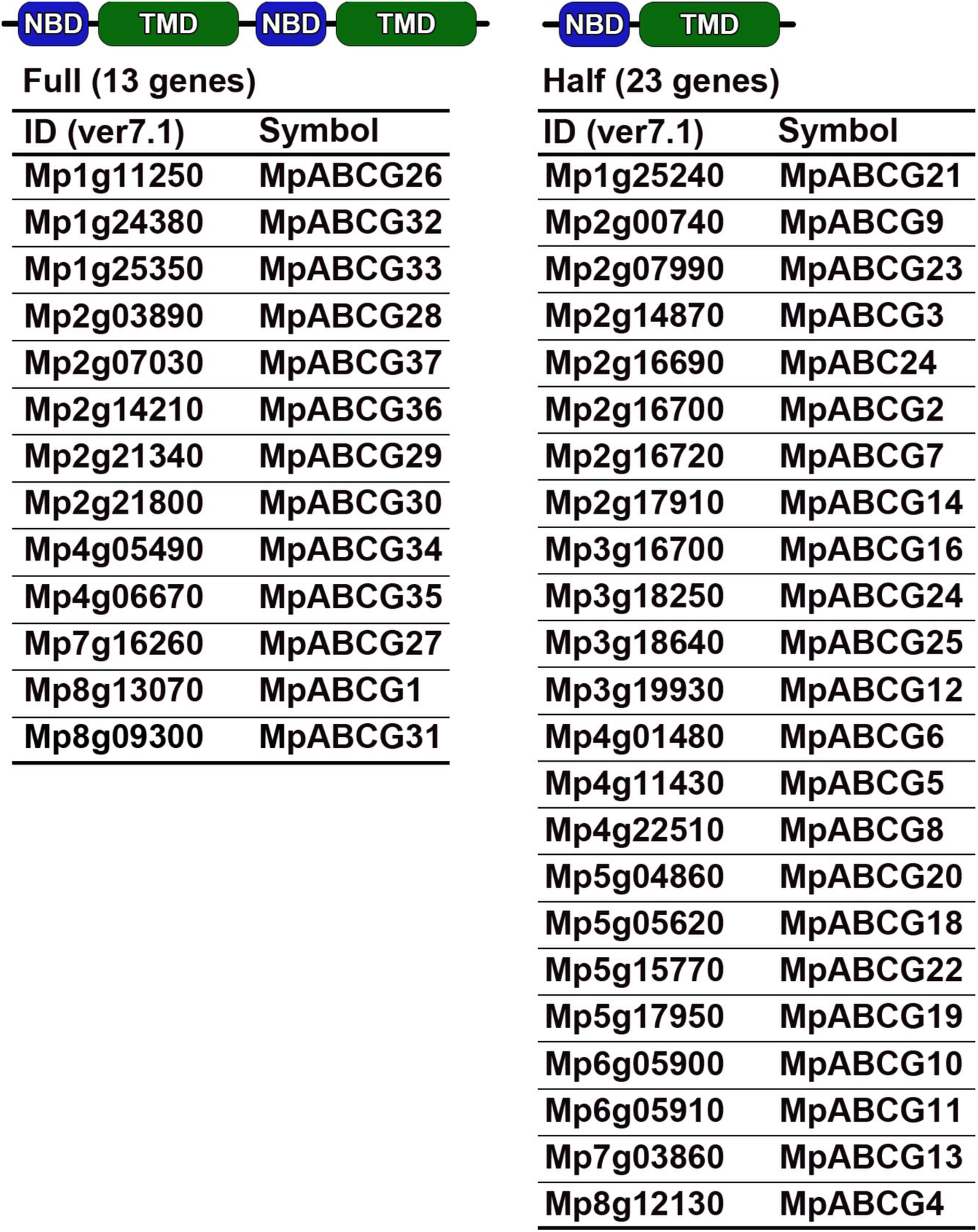
ABCG proteins in *Marchantia polymorpha*. Full transporters consist of two nucleotide-binding domains (NBDs) and two sets of transmembrane domains (TMDs), whereas half transporters consist of only one NBD and a single set of TMDs. In total, 13 and 23 genes were annotated in the *Marchantia* genome database. Gene IDs and symbols are based on MarpolBase (https://marchantia.info/).

**Figure 2.**
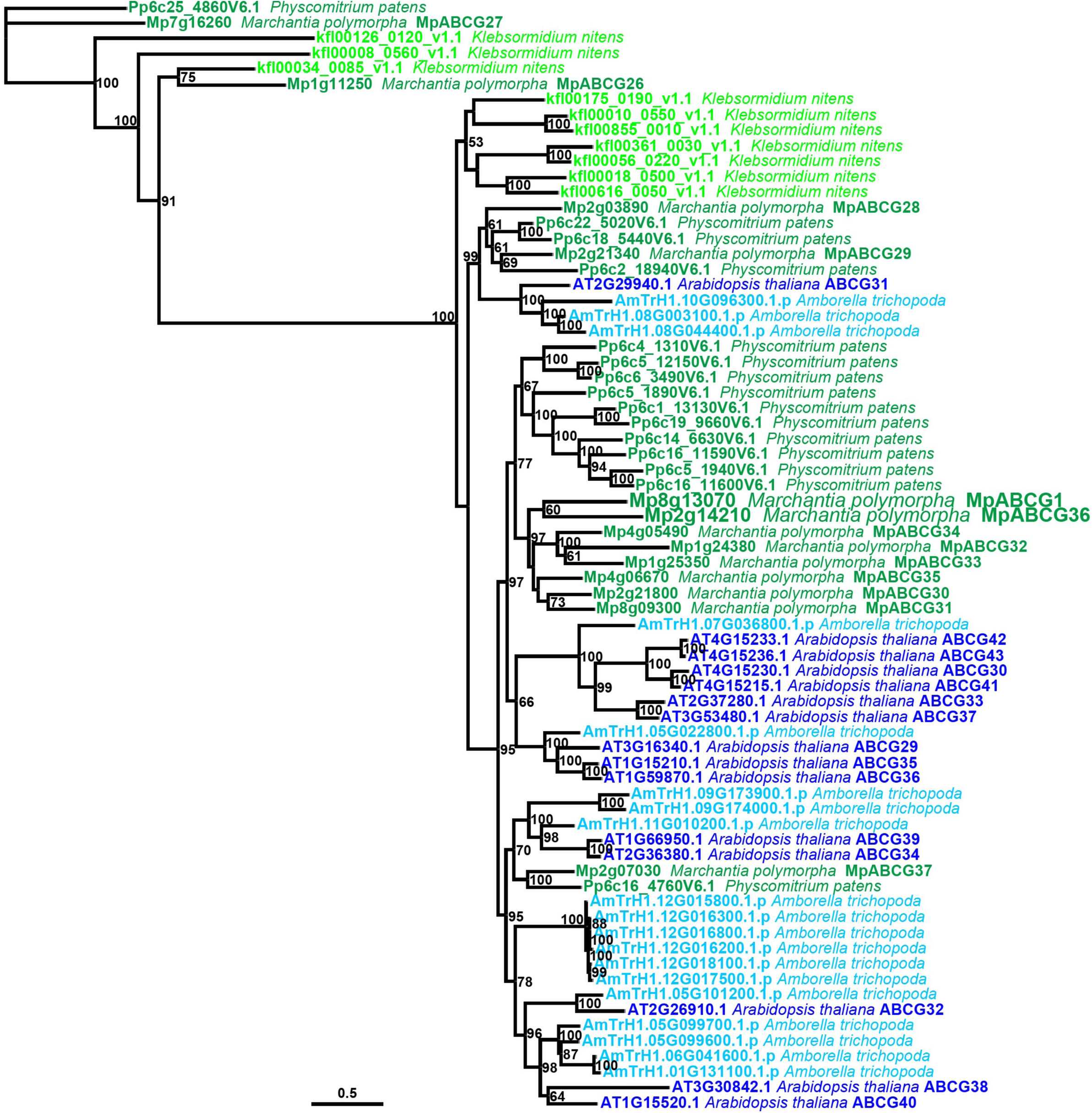
Maximum-likelihood phylogenetic tree of ABCG proteins in plants. Light green, dark green, light blue, and blue operational taxonomic units (OTUs) indicate a charophyte green alga (*Klebsormidium nitens*), bryophytes (liverwort *M. polymorpha* and moss *Physcomitrium patens*), a basal angiosperm (*Amborella trichopoda*), and a eudicot (*Arabidopsis thaliana*), respectively. The branch length is proportional to the estimated number of substitutions per site. Bootstrap probabilities are indicated as percentages on branches with at least 50% support. A more detailed tree was previously constructed by Bowman et al. (2017).

We further examined the expression profiles of ABCG genes using published RNA-sequencing datasets (Kanazawa et al. 2020). Mp*ERF13* is a key transcription factor essential for oil body biogenesis. The gain-of-function (Mp*erf13^GOF^*) and loss-of-function (Mp*erf13-1^ge^*) mutants produce excessive and no oil bodies, respectively. Through the comparison between the three groups using an ANOVA-like test, we identified Mp*ABCG1* and Mp*ABCG36* genes whose expressions were higher than 2 log-fold change in Mp*erf13^GOF^* compared with Tak-1, and lower than −2 log-fold change in Mp*erf13-1^ge^* than Tak-1 (false discovery rate < 0.01) (Figure 3A and B) (Kanazawa et al., 2020). None of the other *ABCG* genes were classified as differentially expressed according to the experimental criteria (Supplementary Figure S1 and S2). Furthermore, mining of single-cell RNA-sequencing data from thallus tissues revealed that the transcripts of both Mp*ABCG1* and Mp*ABCG36* were predominantly enriched in cluster 18, which corresponded to the oil body cell cluster (Supplementary Figure S1 and S2) (Wang et al. 2023). To confirm their tissue distribution and subcellular localization, we generated transgenic *Marchantia* plants expressing mCitrine–MpABCG1 and mCitrine– MpABCG36 under the control of their respective native regulatory elements, including 5′-and 3′-flanking sequences and introns. Both fusion proteins were specifically expressed in oil body cells and localized to the oil body membrane (Figure 3C and D), suggesting that Mp*ERF13* positively regulates the transcription of Mp*ABCG1* and Mp*ABCG36* and that these ABCG proteins function in oil body cells. Next, we generated Mp*abcg1* and Mp*abcg36* mutants via clustered regularly interspaced palindromic repeat (CRISPR)/ Cas9-mediated genome editing and obtained Mp*abcg1*/Mp*abcg36* double mutants by crossing (Figure 4). Mp*abcg1-1*, Mp*abcg1-2*, Mp*abcg36-1*, and Mp*abcg36-2* alleles contained deletions of 8, 1, 175, and 1 bp, respectively, suggesting the loss of function in all cases (Figure 4A and B). Notably, both single (Mp*abcg1-1*, Mp*abcg1-2*, Mp*abcg36-1*, and Mp*abcg36-2*) and double (Mp*abcg1-1*/Mp*abcg36-1*, Mp*abcg1-1*/Mp*abcg36-2*, Mp*abcg1-2*/Mp*abcg36-1*, and Mp*abcg1-2*/Mp*abcg36-2*) mutants exhibited no obvious differences in thallus development compared to the wild-type under the tested experimental conditions (Figure 4C). This phenotype is consistent with previous reports that oil body-deficient mutants (Mp*erf13^ge^*, Mp*c1hdz^ge^*, and Mp*myb02^ge^*) grow normally relative to the wild-type (Kanazawa et al. 2020; Kubo et al. 2024; Romani et al. 2020).

**Figure 3.**
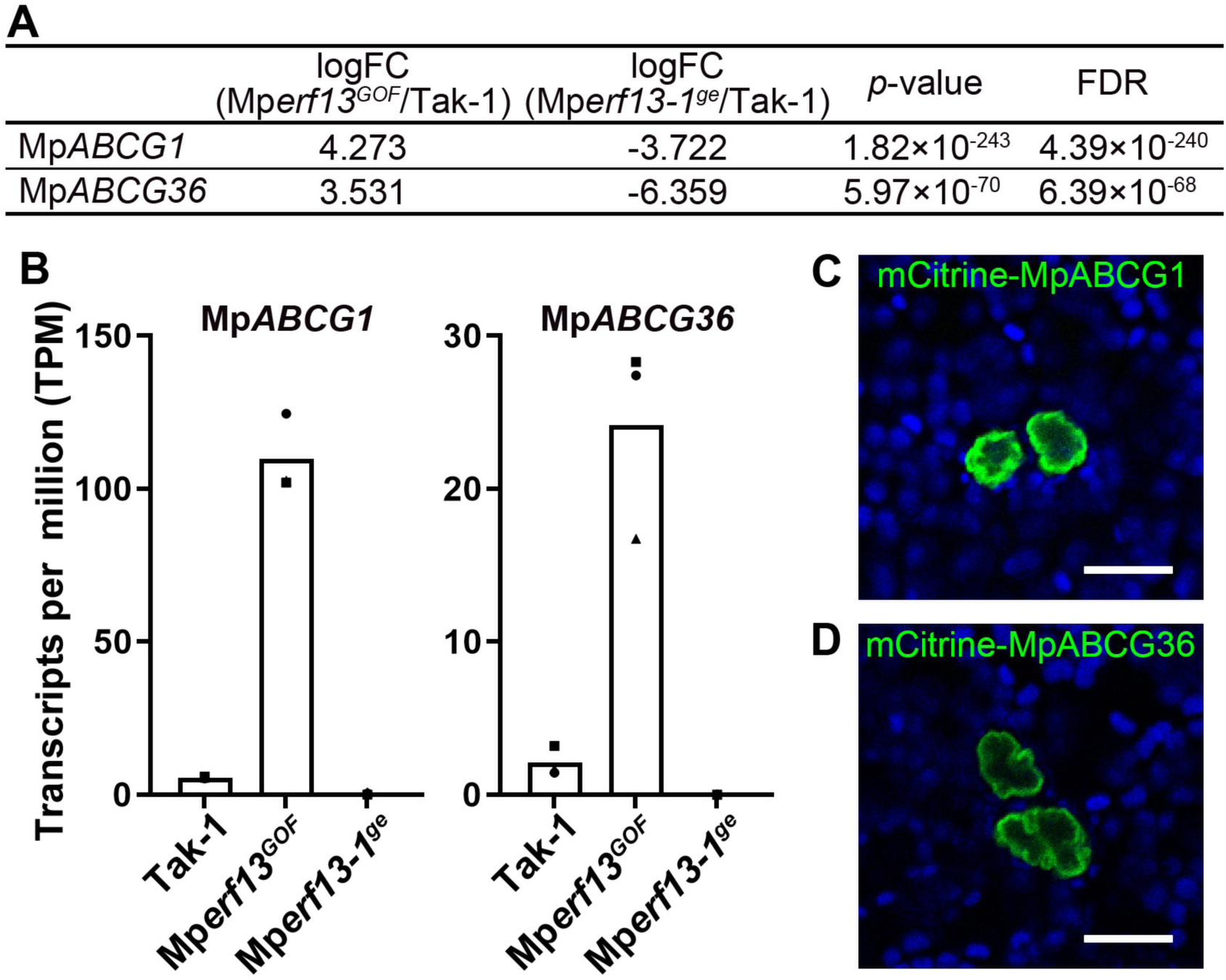
Expression levels of Mp*ABCG1* and Mp*ABCG36* in *M. polymorpha*. (A) Re-analysis of the RNA-sequencing data of Mp*erf13* mutants (Kanazawa et al. 2020). Both Mp*ABCG1* and Mp*ABCG36* exhibited a false discovery rate (FDR) < 0.01 and absolute logFC > 2. (B) Transcripts per million (TPM) values of Mp*ABCG1* and Mp*ABCG36* re-calculated from the RNA-sequencing data (Kanazawa et al. 2020). (C and D) Single confocal images of thallus cells expressing mCitrine–MpABCG1 (C) and mCitrine–MpABCG36 (D) under their native regulatory elements. Green and blue pseudo-colors indicate fluorescence from mCitrine and chlorophyll, respectively. Bars, 20 μm.

**Figure 4.**
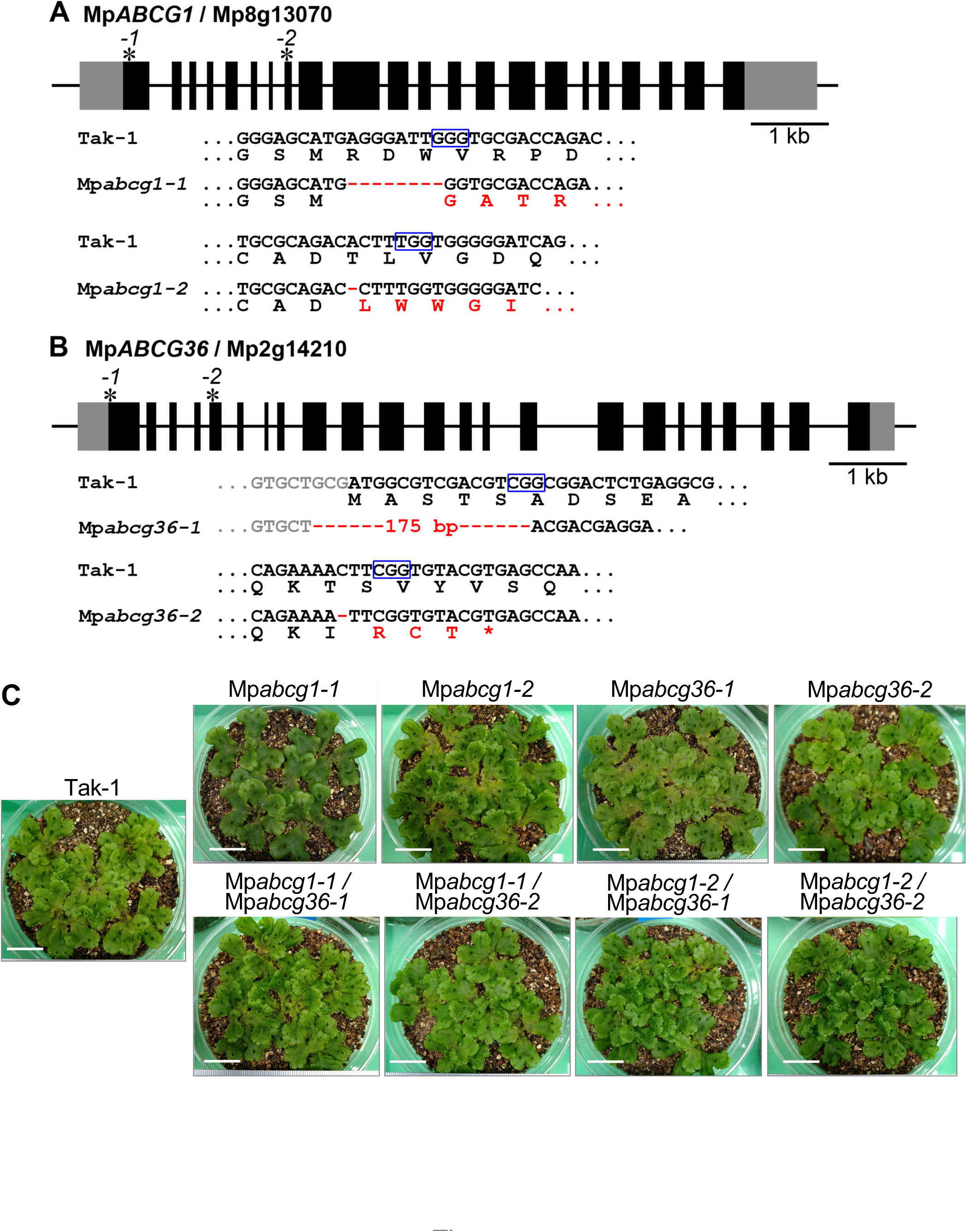
Mp*abcg1* and Mp*abcg36* mutants generated by CRISPR-based genome editing. (A and B) Gene structures of Mp*ABCG1* (A) and Mp*ABCG36* (B). Gray and black boxes represent the untranslated (UTRs) and coding regions, respectively. Asterisks with numbers indicate the mutation sites. PAM and mutated sequences are boxed in blue and highlighted in red, respectively. (C) Six-week-old thalli (two weeks on half-strength B5 agar medium, followed by four weeks on vermiculite) of Tak-1 and Mp*abcg1*, Mp*abcg36*, and Mp*abcg1*/Mp*abcg36* mutants. Genotypes are labeled on each image. Bars, 2 cm.

### Mp*ABCG1* and Mp*ABCG36* mutations exert partially overlapping effects on the accumulation of oil body sesquiterpenoids

Previous microcapillary-based metabolite analysis of oil bodies has demonstrated that sesquiterpenoids, such as thujopsene, chamigrene, and himachalene, accumulate in the oil bodies of *M. polymorpha* (Tanaka et al. 2016). To examine the effects of Mp*ABCG1* and Mp*ABCG36* mutations on the accumulation of these sesquiterpenoids, we conducted gas chromatography (GC)–MS analyses (Figure 5; Supplementary Figures S3 and S4). Among the single mutants, accumulation levels of sesquiterpenes in Mp*abcg1-1* were comparable to those in the wild-type (Tak-1) and those in Mp*abcg36-1* were similar to those in Mp*abcg36-2* (Supplementary Figures S3–S5). Therefore, Mp*abcg1-2* and Mp*abcg36-1* were selected for subsequent analyses. Accumulation of thujopsene in Mp*abcg1-2* significantly decreased to 14.3 ± 1.09% of that in the wild-type, and it was almost undetectable in the Mp*abcg1-2*/Mp*abcg36-1* double mutant (Figure 5B). In Mp*abcg36-1*, thujopsene levels slightly decreased to 75.0 ± 6.09%, although the difference was not statistically significant. Similar trends were observed for β-chamigrene and himachalene, with Mp*abcg1-2* exhibiting significant reductions to 11.2 ± 0.97 and 15.9 ± 0.74%, respectively, compared to Tak-1 (Figure 5B). Moreover, Mp*abcg36-1* exhibited moderate reductions to 75.2 ± 6.99 and 78.2 ± 6.94%, respectively, with the latter showing a statistically significant difference relative to the wild-type. In the double mutant, all three sesquiterpenes were present either below the detection limit or only in trace amounts. Although not all differences were statistically significant, these results collectively suggest that Mp*ABCG1* and Mp*ABCG36* play partially overlapping roles in mediating the accumulation of sesquiterpenoid compounds in the oil bodies of *M. polymorpha*.

**Figure 5.**
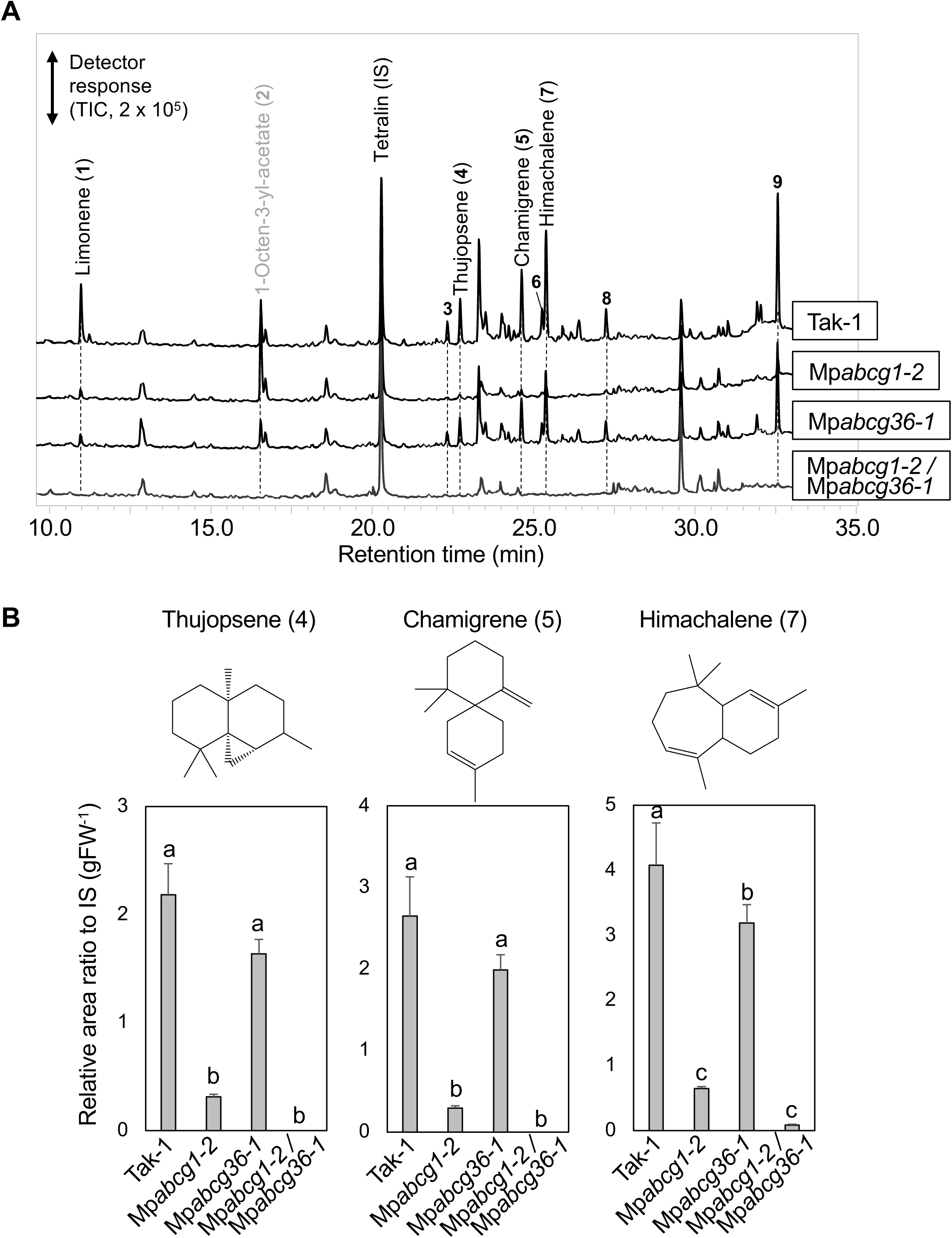
Effects of Mp*abcg1* and Mp*abcg36* mutations on oil body sesquiterpenoid accumulation. (A) Representative gas chromatography–mass spectrometry (GC–MS) chromatograms of Tak-1, Mp*abcg1-2*, Mp*abcg36-1*, and Mp*abcg1-2*/Mp*abcg36-1* thalli. The thalli were cultured for two weeks on half-strength Gamborg’s B5 agar, followed by four weeks on vermiculite. MS profiles of peaks 1–9 are shown in Supplementary Figure S3. Peaks identified with reference compounds are labeled by the compound names. (B) Structures and relative peak areas (mean ± standard error [SE]; n = 3) of thujopsene, chamigrene, and himachalene normalized to the internal standard (IS; tetralin). Different lowercase letters indicate the significant differences (Tukey’s test; P < 0.05).

### Mp*abcg1*, but not Mp*abcg36*, affects the accumulation of the bisbibenzyls, marchantins

In addition to sesquiterpenoids, oil bodies of *M. polymorpha* contain characteristic bisbibenzyl compounds, including marchantin C and its derivatives, which are potentially biosynthesized from the bibenzyl precursor, lunularic acid (Friederich et al. 1999a; Tanaka et al. 2016). To quantify the accumulation of these compounds in the Mp*abcg1* and Mp*abcg36* mutants, we conducted liquid chromatography (LC)–MS/MS analyses (Figure 6; Supplementary Figure S5). Accumulation levels of marchantin C were significantly reduced to 21.7 ± 0.32 and 19.5 ± 1.69% of those in the wild-type (Tak-1) in the Mp*abcg1-2* and Mp*abcg1-2*/Mp*abcg36-1* mutants, respectively, but remained comparable to those in the wild-type in the Mp*abcg36-1* and Mp*abcg36-2* mutants (Figure 6B; Supplementary Figure S5). Similar patterns were observed for marchantin A, whose accumulation levels significantly decreased to 13.7 ± 3.65 and 5.77 ± 1.38% in Mp*abcg1-2* and Mp*abcg1-2*/Mp*abcg36-1*, respectively, but were unaffected in Mp*abcg36-1* and Mp*abcg36-2* (Figure 6B; Supplementary Figure S5). These results suggest that Mp*ABCG1*, but not Mp*ABCG36*, plays a predominant role in the accumulation of bisbibenzyl-type compounds, such as marchantins, in liverwort oil bodies.

**Figure 6.**
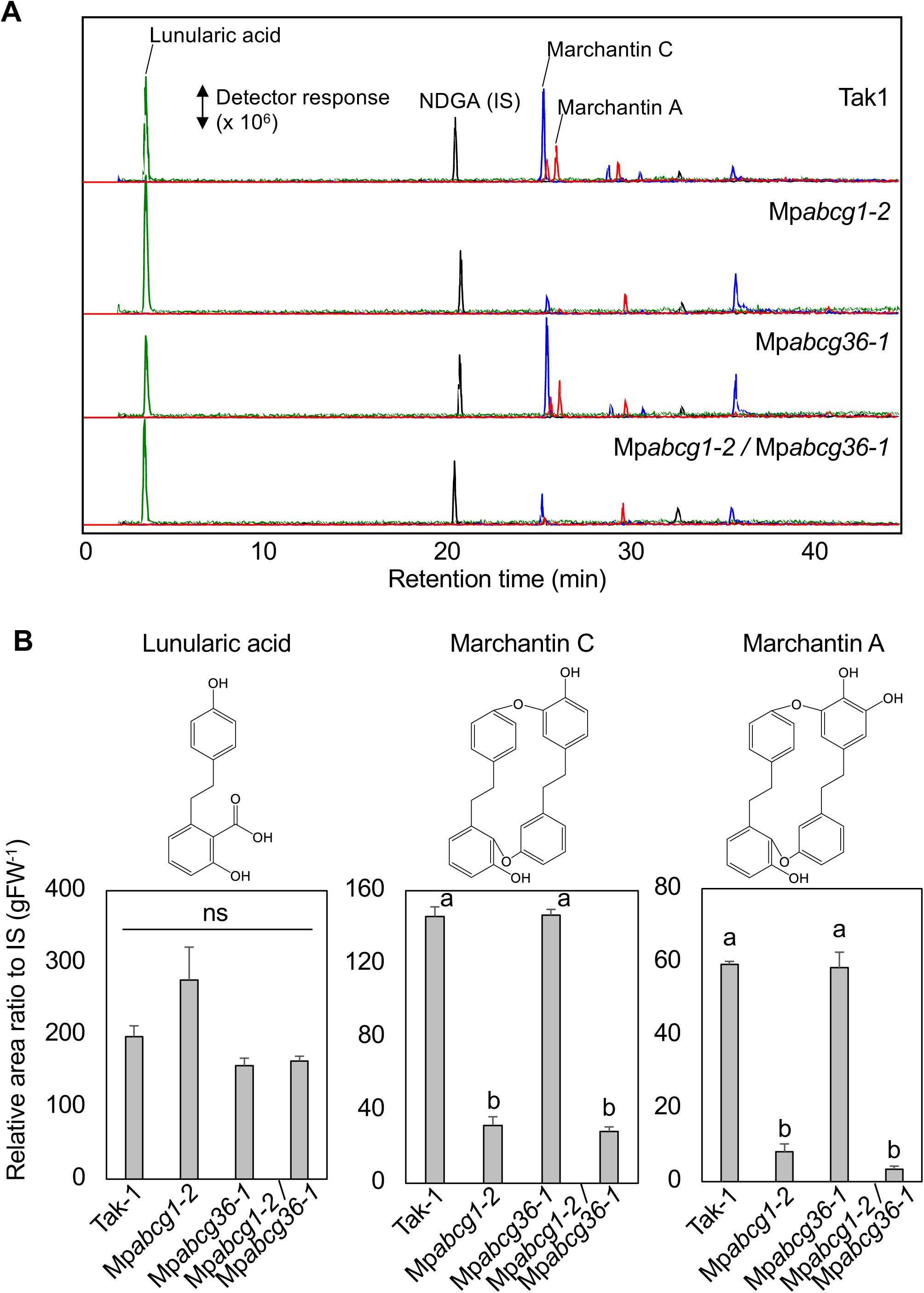
Predominant effect of the Mp*abcg1-2* mutation on marchantin accumulation. (A) Representative liquid chromatography (LC)–MS/MS chromatograms of Tak-1, Mp*abcg1-2*, Mp*abcg36-1*, and Mp*abcg1-2*/Mp*abcg36-1*. Extracted ion chromatograms correspond to lunularic acid (m/z 258.1 ± 0.5; green), nordihydroguaiaretic acid (NDGA; IS; m/z 302.2 ± 0.5; black), marchantin C (m/z 424.2 ± 0.5; blue), and marchantin A (m/z 440.2 ± 0.5; red). (B) Structures and relative peak areas (mean ± SE; n = 3) of lunularic acid, marchantin C, and marchantin A normalized to IS. Different lowercase letters indicate the significant differences (Tukey’s test; P < 0.05).

Notably, the accumulation of lunularic acid, which has not been previously detected in the oil bodies of *M. polymorpha* thalli (Tanaka et al. 2016), showed no significant difference between the Mp*abcg1* and Mp*abcg36* mutants and Tak-1 wild-type (Figure 6B; Supplementary Figure S5).

### Oil bodies in the Mp*abcg1-2* mutant are almost devoid of SMs, as revealed by BODIPY 493/503 staining

Accumulation of sesquiterpenoids and marchantins was reduced in the Mp*abcg1* and Mp*abcg36* mutants (Figures 5 and 6), and their physiological effects were subsequently analyzed. BODIPY 493/503 (hereafter BODIPY), which exhibits a high affinity for lipophilic compounds, is a powerful tool to detect and visualize the oil bodies of *M. polymorpha* (Gutsche et al. 2024; Kanazawa et al. 2020; Kanazawa et al. 2022). The numbers of BODIPY-stained oil bodies per unit area of thalli were 48.3 ± 3.88 and 26.3 ± 5.05 in Mp*abcg1-2* and Mp*abcg1-2*/Mp*abcg36-1*, respectively, with both showing a statistically significant decrease compared to that in Tak-1 (Figure 7A and B).

**Figure 7.**
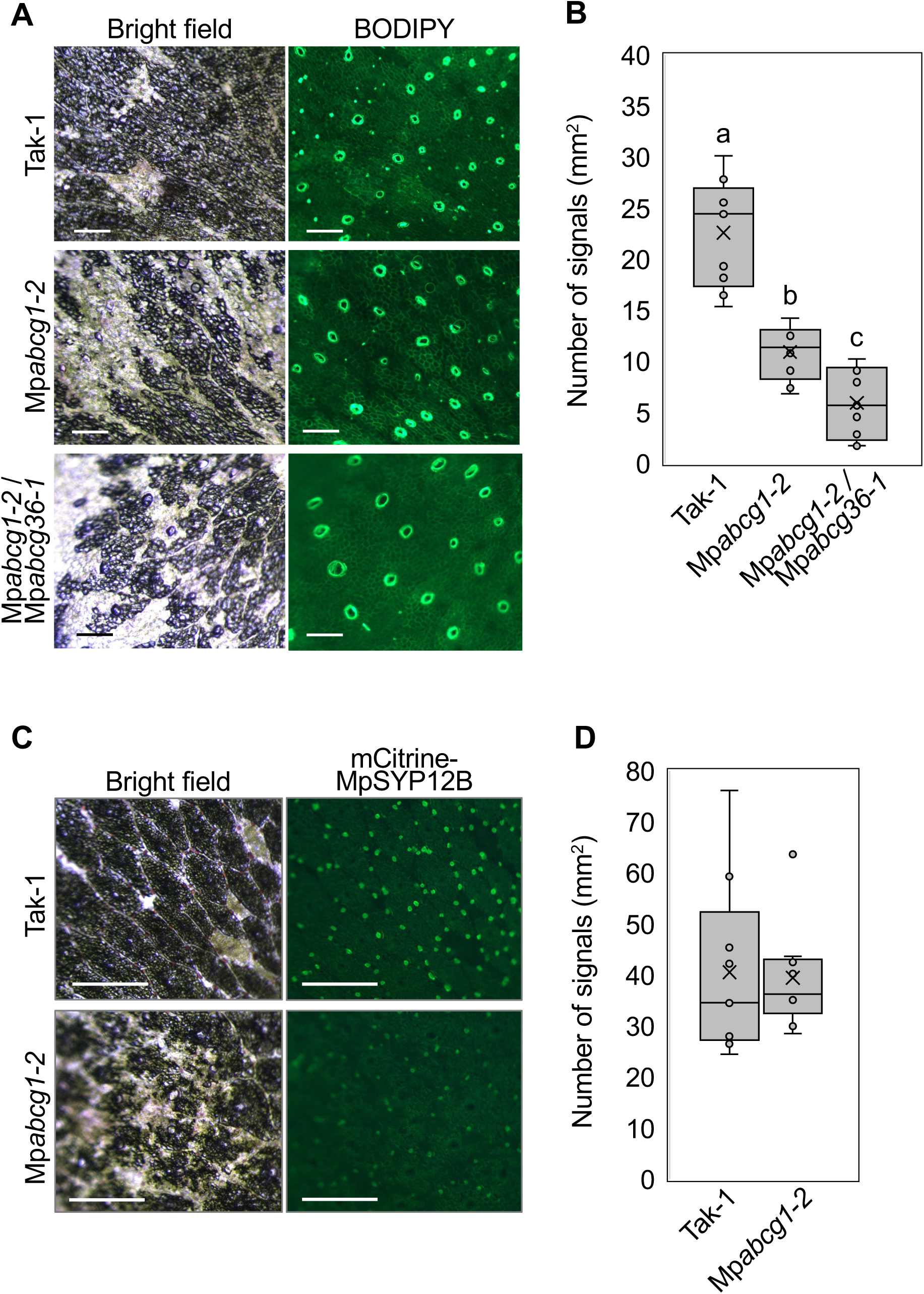
Effect of the Mp*abcg1-2* mutation on the number of BODIPY 493/503-stained oil bodies in *M. polymorpha* thalli. (A) Bright-field and fluorescent images of the BODIPY-stained thalli of Tak-1, Mp*abcg1-2*, and Mp*abcg1-2*/Mp*abcg36-1*. Fluorescence from ring-like structures indicates air pores. Bars, 250 μm. (B) Number of oil bodies visualized by BODIPY per unit area (mm²; n = 9). Bars indicate the mean ± SE. Different lowercase letters indicate the significant differences (Tukey’s test; P < 0.05). (C) Bright-field and fluorescent images of the mCitrine–MpSYP12B-expressing thalli of Tak-1 and Mp*abcg1-2*. Red arrows indicate oil bodies. Bars, 250 μm. (D) Number of oil bodies visualized by mCitrine–MpSYP12B per unit area (mm²). Bars indicate the mean ± SE.

We also examined the oil body membrane marker, mCitrine-MpSYP12B, in Tak-1 and Mp*abcg1-2* via fluorescence stereomicroscopy. mCitrine-MpSYP12B predominantly localizes to the limiting membrane of the oil bodies of *M. polymorpha* (Kanazawa et al. 2020). Fluorescence signals of mCitrine-MpSYP12B were detected in both the wild-type and Mp*abcg1-2* plants (Figure 7C). The number of mCitrine-MpSYP12B-labeled oil bodies per unit area of thalli showed no significant difference (Figure 7D), indicating that oil bodies were normally formed in both the wild-type and Mp*abcg1* mutant. However, Mp*abcg* mutation affected the accumulation of secondary metabolites in oil bodies.

## Discussion

In this study, we demonstrated that ABCG proteins MpABCG1 and MpABCG36 were involved in the accumulation and potential transport of secondary metabolites (SMs) and/or their precursors to liverwort oil bodies and proposed a model for SM accumulation in the oil bodies of *M. polymorpha* (Figure 8). Liverworts produce and accumulate various SMs (He et al. 2013; Kanazawa et al. 2020; Romani et al. 2020). Oil bodies of *M. polymorpha* accumulate many SMs, including sesquiterpenoids (thujopsene, chamigrene, and himachalene) and bisbibenzyls (marchantin C and marchantin A), as revealed via single-cell extraction using microcapillaries, followed by direct MS analysis (Tanaka et al. 2016). However, the mechanisms underlying SM accumulation remain unclear.

**Figure 8.**
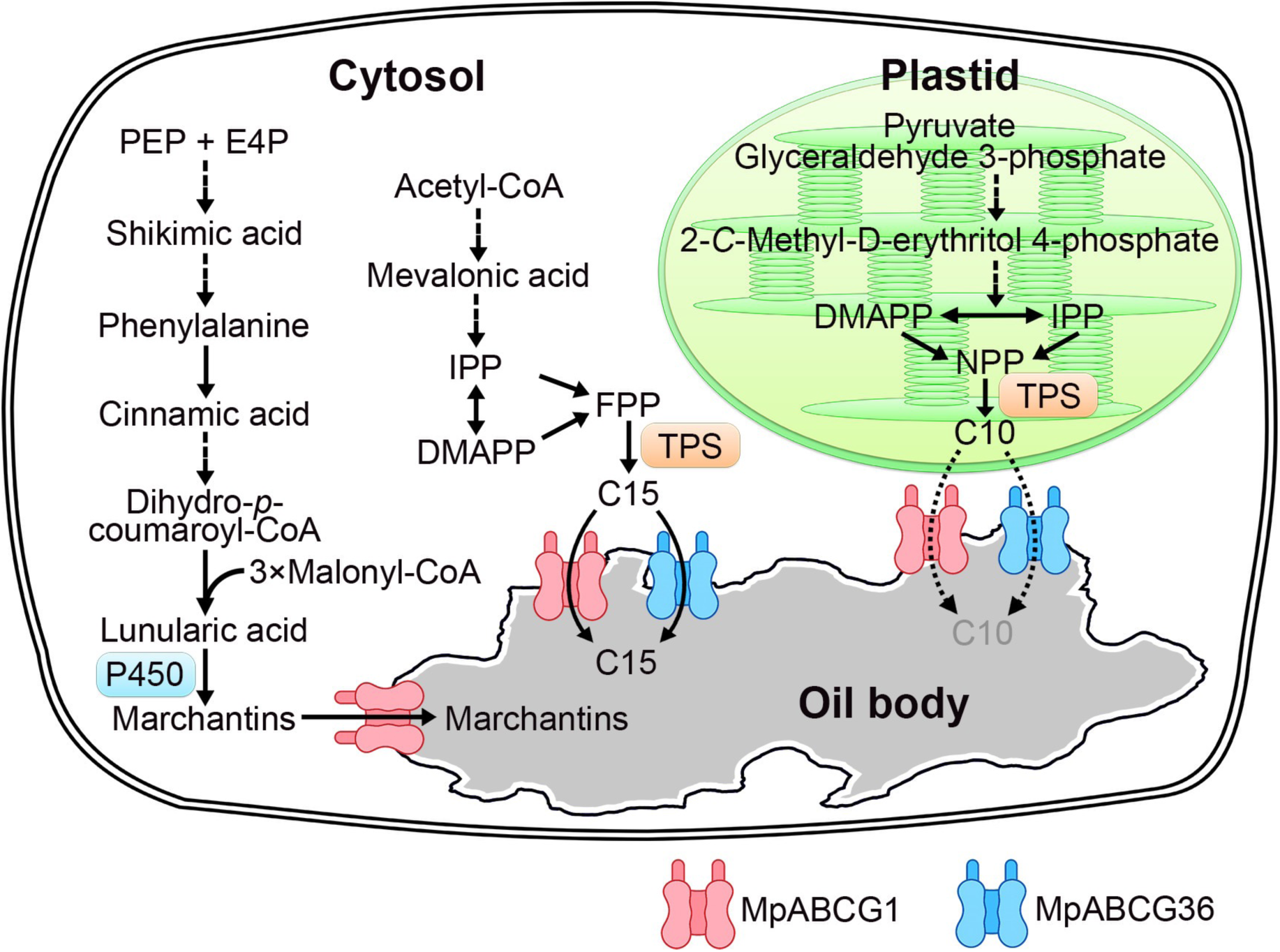
Proposed model of sesquiterpenoid and marchantin accumulation in the oil bodies of *M. polymorpha*. Schematic biosynthetic pathways of sesquiterpenes (C15), marchantins (bisbibenzyls), and monoterpenes (C10) are illustrated primarily based on the existing knowledge on angiosperms, as the corresponding pathways in *M. polymorpha* have not yet been fully elucidated. Both mCitrine–MpABCG1 and mCitrine–MpABCG36 localize specifically to the oil body membrane. MpABCG1 and MpABCG36 play overlapping roles in sesquiterpene accumulation in oil bodies, whereas MpABCG1 plays a predominant role in marchantin accumulation. Although Mp*abcg1*/Mp*abcg36* mutations affect the biosynthesis and/or accumulation of monoterpenes (C10), their oil body distribution remains unclear. Abbreviations: PEP, phosphoenolpyruvic acid; E4P, erythrose 4-phosphate; P450, cytochrome P450; IPP, isopentenyl pyrophosphate; DMAPP, dimethylallyl pyrophosphate; NPP, neryl pyrophosphate; FPP, farnesyl pyrophosphate; TPS, terpene synthase; C10, monoterpenoids; C15, sesquiterpenoids.

This study first focused on MpABCG1, which was previously reported to localize to the oil body membrane in *M. polymorpha* (Kanazawa et al. 2020). Re-analysis of the RNA-sequencing data of Mp*erf13* mutants and confocal microscopic observation of fluorescent-protein fusions revealed that both MpABCG1 and MpABCG36 function at the oil body membrane (Figures 1–3). Levels of oil body sesquiterpenes (thujopsene, chamigrene, and himachalene) were decreased in the Mp*abcg1* and Mp*abcg36* mutants generated via CRISPR–Cas9-mediated genome editing. Moreover, the accumulation of bisbibenzyls (marchantin C and marchantin A) was reduced in Mp*abcg1*, but not in Mp*abcg36* (Figures 4–6). Although SM accumulation was reduced, the number of mCitrine-MpSYP12B-labeled oil bodies in the Mp*abcg1-2* mutant was comparable to that in the wild-type (Figure 7), suggesting that MpABCG1 and MpABCG36 are essential for SM accumulation in oil bodies, but not for oil body formation.

Mp*ABCG1* was initially identified as a differentially expressed gene via RNA-sequencing analysis of Mp*erf13* mutants, and mCitrine-MpABCG1 was reported to be specifically expressed in oil body cells and localized to the oil body membrane (Kanazawa et al. 2020). In this study, we generated two mutant alleles, Mp*abcg1-1* and Mp*abcg1-2*, the latter of which showed the reduced accumulation of sesquiterpenes and marchantins. Considering the completely different biosynthetic pathways of sesquiterpenes and marchantins, MpABCG1 is possibly involved in the accumulation of diverse compounds. Although Mp*abcg1-1* was predicted to be a frameshift mutation introducing a premature stop codon, it did not affect the accumulation of sesquiterpenes and marchantins (Figures 5 and 6; Supplementary Figures S4 and S5), probably because an unpredicted in-frame protein was still translated. Forestier et al. (2025) independently generated loss-of-function Mp*abcg1* mutant lines and demonstrated a marked reduction in sesquiterpene accumulation in these lines. These findings suggest that MpABCG1 plays an essential role in sesquiterpene accumulation in oil bodies.

In this study, Mp*abcg1-2*/Mp*abcg36-1* mutant exhibited the reduced accumulation of three sesquiterpenes (thujopsene, chamigrene, and himachalene), sesquiterpene oxides, monoterpenes (e.g., limonene), bisbibenzyls, and an oxylipin-related compound, 1-octen-3-yl acetate (Figure 5). This suggests that MpABCG1 and MpABCG36 play pleiotropic roles in the biosynthesis and accumulation of structurally diverse compounds. However, whether compounds other than sesquiterpenes and marchantins also accumulate in oil bodies remains unclear, warranting further investigation. Existing knowledge on angiosperms suggests that sesquiterpenoids, marchantins, and monoterpenes are biosynthesized in the cytosol or plastids (Figure 8) (Vranová et al. 2013), through the pathways involving several biosynthetic enzymes and cofactors, such as NADPH and acetyl-CoA (Chapple 1998; Kihara et al. 2014). However, their functional subcellular distribution in *M*. *polymorpha* has not yet been fully characterized. In addition to the systematic analyses of sesquiterpene biosynthetic enzyme expression and localization, as conducted by Forestier et al. (2025), future studies should perform ABCG transporter assays, as performed by Adebesin et al. (2017), using liposomes and/or isolated membranes to identify the correct substrate(s).

Oil bodies contain specific bisbibenzyl compounds, including marchantin C and its derivatives (Tanaka et al. 2016). Our study indicated that MpABCG1 was a major contributor to the accumulation of both marchantins C and A in oil bodies. P450 enzymes are necessary for the conversion of marchantin C to marchantin A (Friederich et al. 1999b), and most canonical P450 enzymes function at the ER (Chapple 1998); therefore, marchantins C and A are possibly biosynthesized on the cytosolic surface of the ER and subsequently transported to the lumen of oil bodies. As reported for volatile organic compounds (Widhalm et al. 2015), inhibition of marchantin sequestration may lead to the excessive accumulation of highly hydrophobic marchantins in the cytosol, thereby impairing the endomembrane integrity. In this study, marchantin levels were lower in the Mp*abcg1-2* mutant than in the wild-type, and plant growth appeared normal under the tested experimental conditions (Figures 4C and 6). These finding suggest the presence of a feedback control system and/or homeostatic mechanism that regulates the metabolic flux and marchantin accumulation. Lunularic acid levels did not differ significantly between Tak-1 and Mp*abcg1-2*, and similar metabolic profiles have been reported in Mp*myb02* (Kubo et al. 2018). Unlike Mp*abcg1-2*, Mp*myb02* undergoes oil body cell differentiation, but not oil body formation (Kubo et al. 2024; Wang et al. 2023). Comparative analyses of these mutants will provide deeper insights into the feedback control system and/or homeostatic mechanism preventing cellular toxicity caused by excessive marchantin accumulation. Importantly, our findings highlight the compartmentalization of SMs at high concentrations by organelles that have evolved in a lineage-specific manner.

## Materials and Methods

### Phylogenetic analysis

Protein sequences were collected from Phytozome 13 and the following genome portal sites: *Amborella trichopoda* v1.0, *Arabidopsis thaliana* TAIR11, *Klebsormidium nitens* NIES-2285 v1.1 (http://www.plantmorphogenesis.bio.titech.ac.jp/~algae_genome_project/klebsormidium/klebsormidium_blast.html), *M. polymorpha* v5.1, and *Physcomitrium patens* v3.3. Multiple sequence alignments were performed using MUSCLE version 3.8.31 with default parameters (Edgar 2004a, b). Maximum-likelihood phylogenetic analyses were conducted using PhyML 3.0 under LG model (Guindon et al. 2010). Bootstrap support values were estimated from 1,000 replicates.

### RNA-sequencing re-analysis

RNA-sequencing data (accession no. DRA009193) were used for re-analysis (Kanazawa et al. 2020). Transcript abundance was quantified using kallisto (version 0.43.1) with default parameters and the primary transcript dataset from *Marchantia* genome version 3.1 (Bowman et al. 2017; Bray et al. 2016). Single-cell RNA-sequencing data were mined using Single Cell Browser (http://wanglab.sippe.ac.cn/Marchantia-census/) (Wang et al. 2023).

### DNA constructs

All oligonucleotides and DNA constructs used in this study are listed in Table 1. To construct mCitrine–MpABCG36, a 14.5-kb genomic fragment of Mp*ABCG36* containing the 5′- and 3′-flanking sequences was amplified by PCR from Tak-1 genomic DNA, and the amplified product was subcloned into pENTR/D-TOPO (Thermo Fisher Scientific, Waltham MA, USA), according to the manufacturer’s instructions. The coding sequence of mCitrine was subsequently inserted upstream of the initiation codon of Mp*ABCG36* using the In-Fusion HD Cloning Kit (Takara, Shiga, Japan), according to the manufacturer’s instructions. The resultant construct was introduced into the pMpGWB301 Gateway binary vector (Ishizaki et al. 2015) using the Gateway LR Clonase II Enzyme Mix (Thermo Fisher Scientific).

**Table 1.**
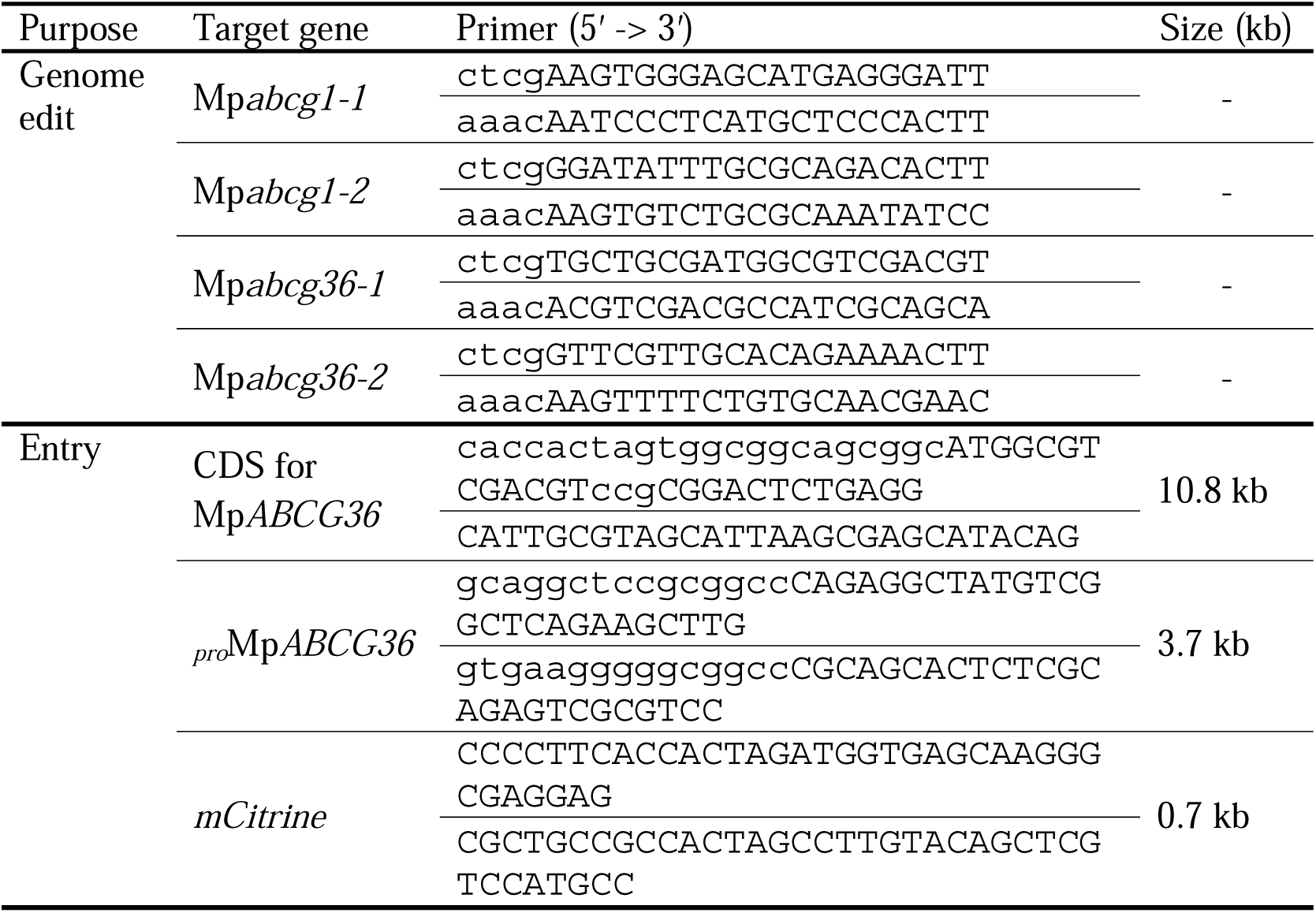
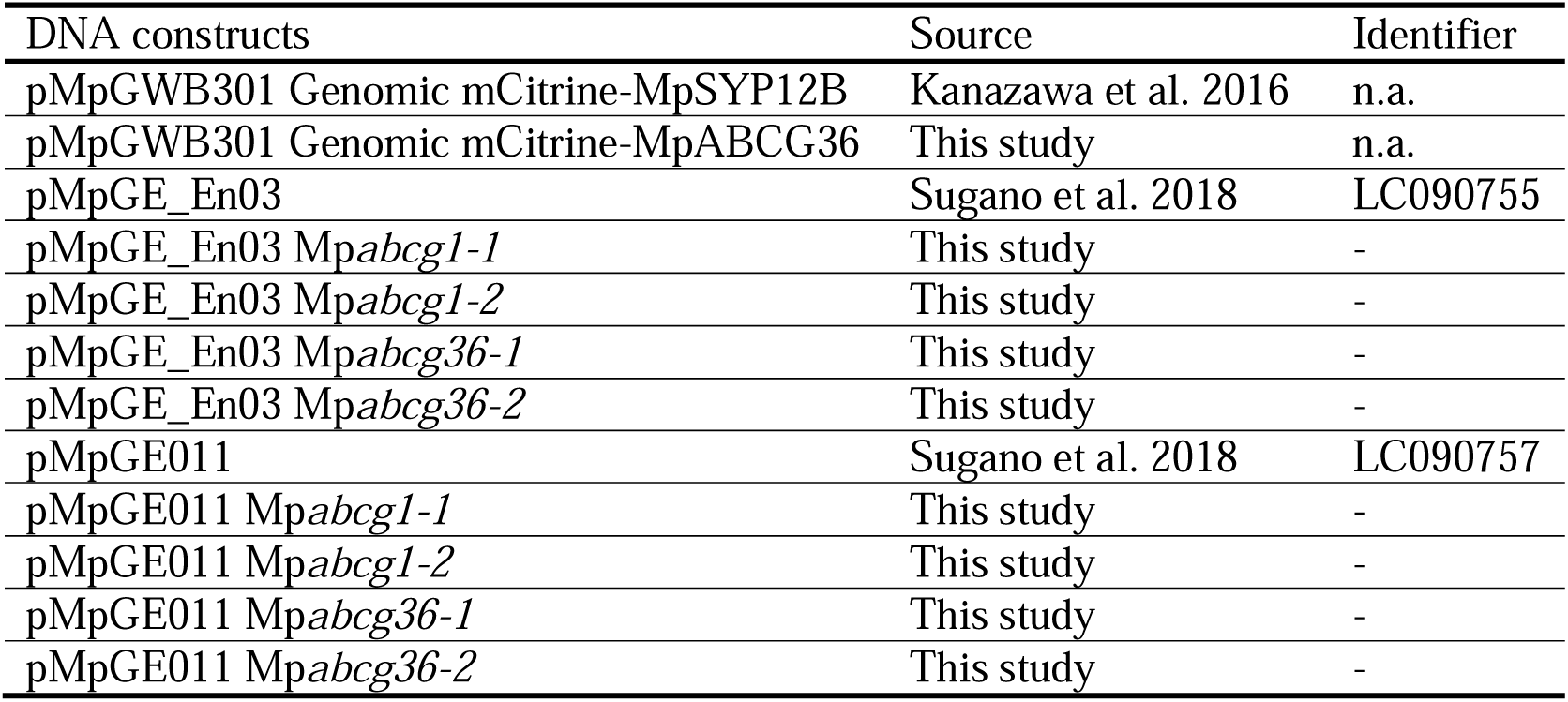
List of primers and DNA constructs used in this study.

To construct genome-editing vectors, double-stranded DNA fragments were inserted into the BsaI site of pMpGE_En03 (Sugano et al. 2018). The resultant constructs were introduced into the pMpGE011 Gateway binary vector (Sugano et al. 2018) using the Gateway LR Clonase II Enzyme Mix, following the manufacturer’s instructions.

### Plant growth and transformation

All strains used in this study are listed in Table 2. Male and female accessions of *M. polymorpha* Takaragaike-1 (Tak-1) and Takaragaike-2 (Tak-2), respectively, were used for analyses (Ishizaki et al. 2008). Growth conditions and transformation methods were as previously described (Kanazawa et al. 2020; Kubota et al. 2013; Takizawa et al. 2021). Gemmae were aseptically cultured on half-strength Gamborg’s B5 medium (Fujifilm Wako Pure Chemicals, Osaka, Japan) containing 1% agar in plastic Petri dishes (diameter: 90 mm; depth: 20 mm; Sansei Medical, Kyoto, Japan) at 22 °C under continuous fluorescent light (36–61 µmol photons m[² s[¹). After cultivation on agar plates for two weeks, the thalli were transplanted onto vermiculite (thickness: 5 cm) soaked in 2,000-fold diluted Hyponex nutrient solution (Hyponex Japan, Osaka, Japan). The thalli were further cultured on vermiculite at 22 °C under continuous fluorescent light in a growth chamber. The vermiculite was kept moist via regular watering, and no additional nutrients were supplied after initial application of the diluted Hyponex solution. Far-red light was supplemented to induce the transition from vegetative to reproductive growth. Mutants generated via genome editing were crossed with the wild-type accessions to remove the Cas9 cassette.

**Table 2.**
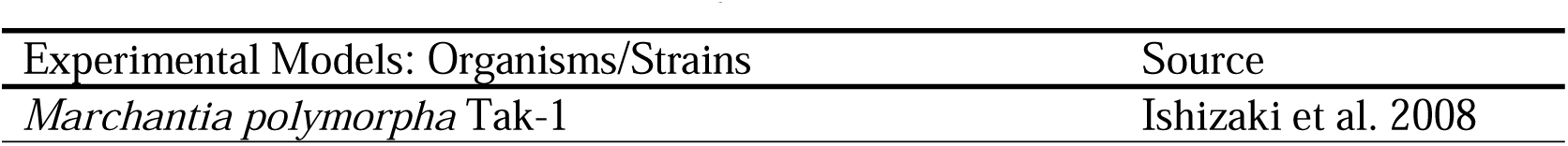
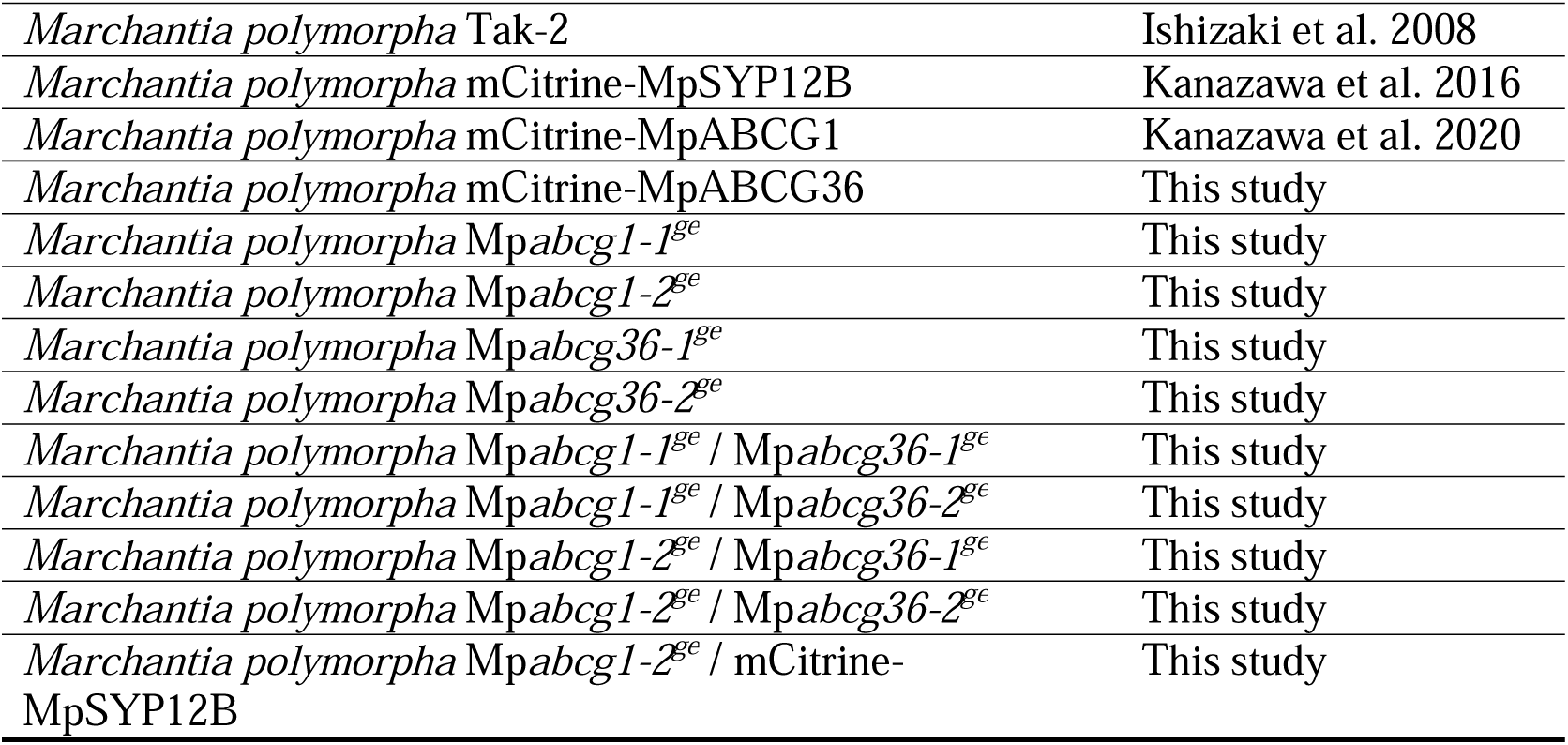
List of strains used in this study.

### GC–MS analysis of volatile compounds

Thalli grown for four weeks on vermiculite after transplanting from agar plates were used for volatile analysis, as previously described (Takizawa et al. 2021). The thalli (approximately 100 mg fresh weight) were placed in a 2-mL screw-cap tube (Sarstedt, Nümbrecht, Germany) and snap-frozen in liquid nitrogen with eight stainless steel beads (diameter: 3 mm). After the addition of 0.4 mL methyl *tert*-butyl ether (Fujifilm Wako Pure Chemicals, Osaka, Japan) containing 2.0 µg mL[¹ tetralin as an internal standard (IS), the thalli were vigorously lysed using a bead cell disrupter (Micro Smash MS-100R; Tomy, Tokyo, Japan) at 3,500 rpm for 1 min. The mixture was centrifuged at 15,000 rpm for 3 min at 25 °C, and the upper organic phase was directly subjected to GC–MS (Tanaka et al. 2016). A GC–MS system (QP-5050; Shimadzu, Kyoto, Japan) equipped with the DB-WAX column (length × inner diameter: 30 m × 0.25 mm; film thickness: 0.25 µm; Restek, Bellefonte, PA, USA) was used. The column temperature was set as follows: 80 °C for 5 min, increased to 240 °C at 10 °C min[¹, and held for 5 min. Helium was used as the carrier gas at a flow rate of 40 cm s[¹. Injector and interface temperatures were set at 200 °C. Splitless injections were performed with a sampling time of 1 min. The mass spectrometer was operated in the electron impact mode with an ionization energy of 70 eV. The relative area ratio of each peak to that of tetralin (IS) was used to quantify each volatile compound. Thujopsene, chamigrene, and himachalene were identified by comparing their MS spectra and retention times with those of authentic standards kindly provided by Dr. H. Kenmoku and Dr. Y. Asakawa (Tokushima Bunri University, Tokushima, Japan). Limonene and 1-octen-3-yl acetate were purchased from Fujifilm Wako Pure Chemicals. Other compounds were tentatively identified by comparing their MS spectra with those in the National Institute of Standards and Technology-17 database (Shimadzu).

### LC–MS/MS analysis of marchantins

Marchantin levels were estimated as previously described (Tanaka et al. 2016). Apical portion of the thalli containing the primordia (approximately 10 mg) was excised with a razor blade, placed in a 2.0-mL screw-cap microtube with seven stainless steel beads (diameter: 3 mm), snap-frozen in liquid nitrogen, and stored at –80 °C until use. To each sample, 400 µL methanol containing 20.0 µg mL[¹ nordihydroguaiaretic acid (Fujifilm Wako Pure Chemicals) as IS was added, followed by two rounds of homogenization at 3,500 rpm for 1 min using a bead-type cell disruptor (Micro Smash MS-100R; Tomy). After centrifugation at 15,000 × g for 3 min, the supernatant was directly analyzed using the 3200 QTRAP system (AB Sciex, Framingham, MA, USA) equipped with the Prominence UFLC system (Shimadzu).

Marchantins were separated on the Mightysil RP-18 column (150 mm × 2 mm; 5 µm; Kanto Chemical, Tokyo, Japan) with a binary solvent system consisting of H[O/HCOOH (100:0.1 [v/v]; solvent A) and CH[CN/HCOOH (100:0.1 [v/v]; solvent B). The elution program consisted of a linear gradient from 20% solvent B (5 min) to 90% solvent B for 35 min at a flow rate of 0.2 mL min[¹. Detection was performed using a photodiode array detector (SPD-M20A; Shimadzu) and MS/MS with electrospray ionization in the positive-ion mode (ion spray voltage: 3,000 V; temperature: 120 °C; N[as curtain gas [12 arbitrary units] and collision gas set to “high”; collision energy: 6 V; declustering potential: 30 V). The relative amounts of marchantins were calculated from the ratio of the analyte peak area to that of IS.

### Stereomicroscopy

Apical portion of the thalli (1 cm) was excised using a razor blade and examined under a fluorescence stereomicroscope (AF6000 Macro; Leica, Wetzlar, Germany). A GFP3 filter set (excitation: 470/40 nm; emission: 525/50 nm) was used to detect BODIPY and mCitrine fluorescence. For BODIPY staining, the excised thallus piece was immersed in 1 mL of 3.82 µM BODIPY solution in darkness for 15 min.

### Confocal laser-scanning microscopy

Five-day-old thalli were used for observation. The samples were mounted in half-strength Gamborg’s B5 liquid medium and observed using the LSM780 confocal microscope (Carl Zeiss, Germany). Excitation was performed at 488 nm (argon laser) and 561 nm (DPSS 561–10), and emissions were collected between 482 and 659 nm. Spectral unmixing of the obtained images was performed using the ZEN 2.3 SP1 software (Carl Zeiss). Representative confocal microscopy images of 20 biological replicates are shown.

## Abbreviations

ABC: ATP-binding cassette
BODIPY: 4,4-difluoro-1,3,5,7,8-pentamethyl-4-bora-3a,4a-diaza-*s*-indacene
CRISPR: clustered regularly interspaced short palindromic repeat
ER: endoplasmic reticulum
FPP: farnesyl diphosphate
FTPSL: fungal-type terpene synthase-like
GC–MS: gas chromatography–mass spectrometry
gRNA: guide RNA
IPP: isopentenyl diphosphate
LC–MS/MS: liquid chromatography–tandem mass spectrometry
mCitrine: monomeric Citrine
NBD: nucleotide-binding domain
PM: plasma membrane
RT-qPCR: reverse transcription-quantitative PCR
SM: specialized metabolite
TMD: transmembrane domain
TPS: terpene synthase
MVA: mevalonic acid
MEP: 2-*C*-methyl-D-erythritol 4-phosphate

## Acknowledgments

We would like to thank Dr. Takayuki Kohchi (Kyoto Universitym Kyoto, Japan), Dr. Ryuichi Nishihama (Tokyo University of Science, Chiba, Japan), Dr. Kimitsune Ishizaki (Kobe University, Kobe, Japan), Dr. Shigeo S. Sugano (AIST, Tsukuba, Japan), Dr. Yoshinori Asakawa (Tokushima Bunri University, Tokushima, Japan), and Dr. Hiromichi Kenmoku (Tokushima Bunri University, Tokushima, Japan) for sharing the vectors, plants, and specialized metabolite (SM) standard materials used in this study. *Marchantia* culture was supported by the Model Plant Section of the Model Organisms Facility, NIBB Trans-Scale Biology Center (Okazaki, Japan).

## Supporting information

**Supplementary Figure S1. Transcriptional expression profiles of Mp*ABCG1* and Mp*ABCG36*.** (A) Uniform manifold approximation and projection (UMAP) visualization of the cell clusters in *M. polymorpha* reported by Wang et al. (2023). (B) Mp*ABCG1* expression levels in UMAP clusters. (C) Mp*ABCG36* expression levels in UMAP clusters. Both Mp*ABCG1* and Mp*ABCG36* were predominantly plotted in cluster 18 (oil body cell cluster). UMAPs were generated using http://wanglab.sippe.ac.cn/Marchantia-census/ (Wang et al. 2023).

**Supplementary Figure S2. Transcriptional expression profiles of all *ABCG* genes in *M*. *polymorpha*.** Expression levels of all *ABCG* genes are shown as UMAP cluster distributions generated using Single Cell Browser (Wang et al., 2023) and transcripts per million (TPM) values re-calculated from the RNA-sequencing data of Kanazawa et al. (2020). Data for Mp*ABCG1* and Mp*ABCG36* are presented in Figure 3 and Supplementary Figure S1, respectively. Full and half *ABCG* genes are presented in panels (A–K) and (L–AH), respectively.

**Supplementary Figure S3. Mass spectra of compounds 1–9.** For each compound, the most similar match in the National Institute of Standards and Technology (NIST)-17 library is shown, with the similarity score indicated in parentheses.

**Supplementary Figure S4. Levels of sesquiterpenoids in all mutants generated in this study.** Levels of three sesquiterpenoids identified in the extracts of the six-week-old thalli of all mutants generated in this study are shown. The relative peak area of each compound normalized to the internal standard (IS, tetralin) is shown. Data are represented the mean ± SE of three biological replicates. Different lowercase letters indicate the significant differences (one-way analysis of variance [ANOVA], followed by Tukey’s test; P < 0.05).

**Supplementary Figure S5. Levels of three bibenzyls in all mutants generated in this study.** Levels of three bibenzyls identified in the extracts of the six-week-old thalli of all mutants generated in this study are shown. The relative peak area of each compound normalized to IS (nordihydroguaiaretic acid [NDGA]) is shown. Data are represented as the mean ± SE of three biological replicates. Different lowercase letters indicate the significant differences (one-way ANOVA, followed by Tukey’s test; P < 0.05).

