## Supplemental Figures for "ABCG transporters are involved in the accumulation of specialized metabolites in the oil bodies of *Marchantia polymorpha*"

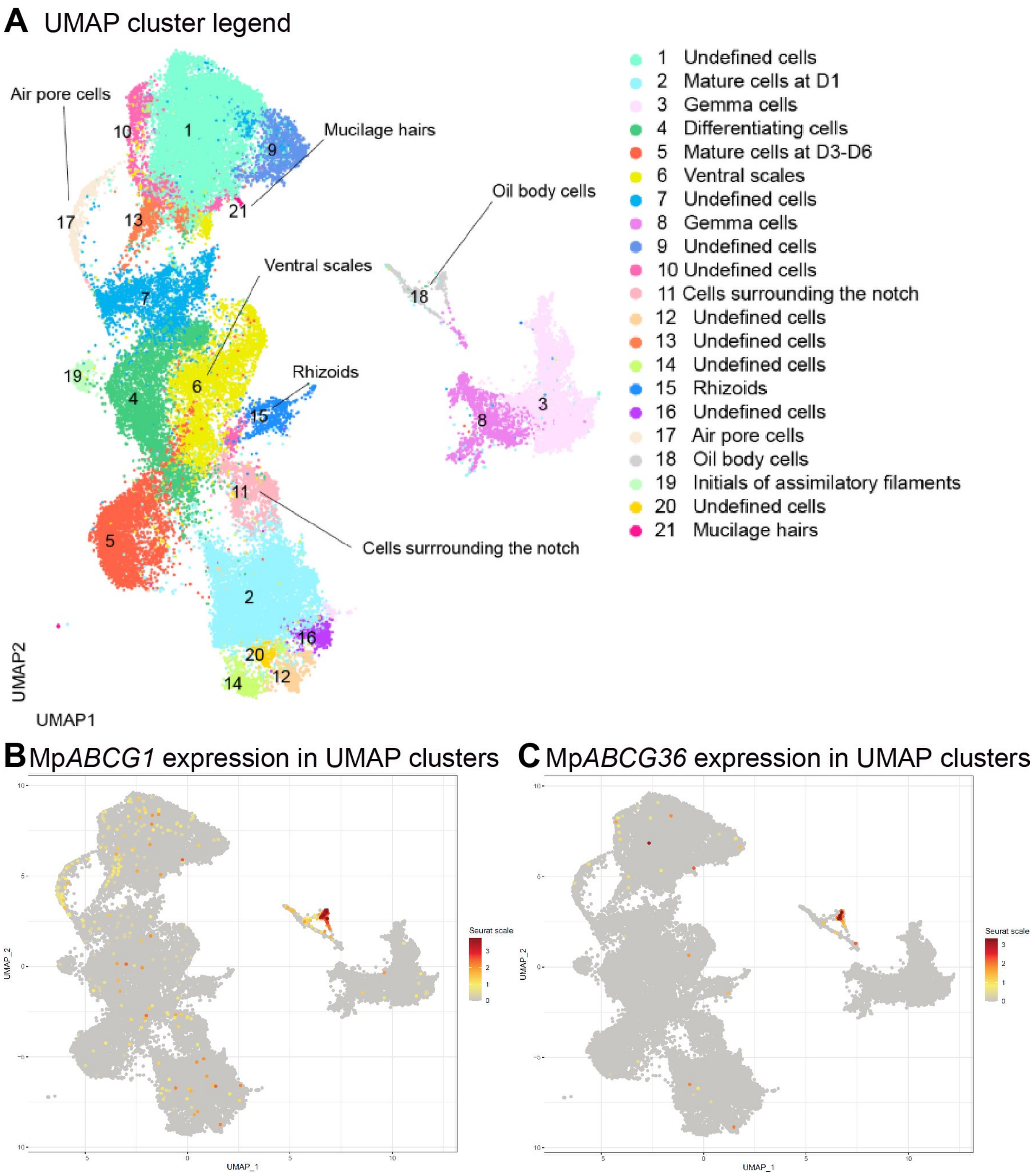


**Supplementary Figure S1. Transcriptional expression profiles of Mp*ABCG1* and Mp*ABCG36*.**

**(A)** Uniform manifold approximation and projection (UMAP) visualization of the cell clusters in *M. polymorpha* reported by Wang et al. (2023). **(B)** Mp*ABCG1* expression levels in UMAP clusters. (C) Mp*ABCG36* expression levels in UMAP clusters. Both Mp*ABCG1* and Mp*ABCG36* were predominantly plotted in cluster 18 (oil body cell cluster). UMAPs were generated using http://wanglab.sippe.ac.cn/Marchantia-census/ (Wang et al. 2023).

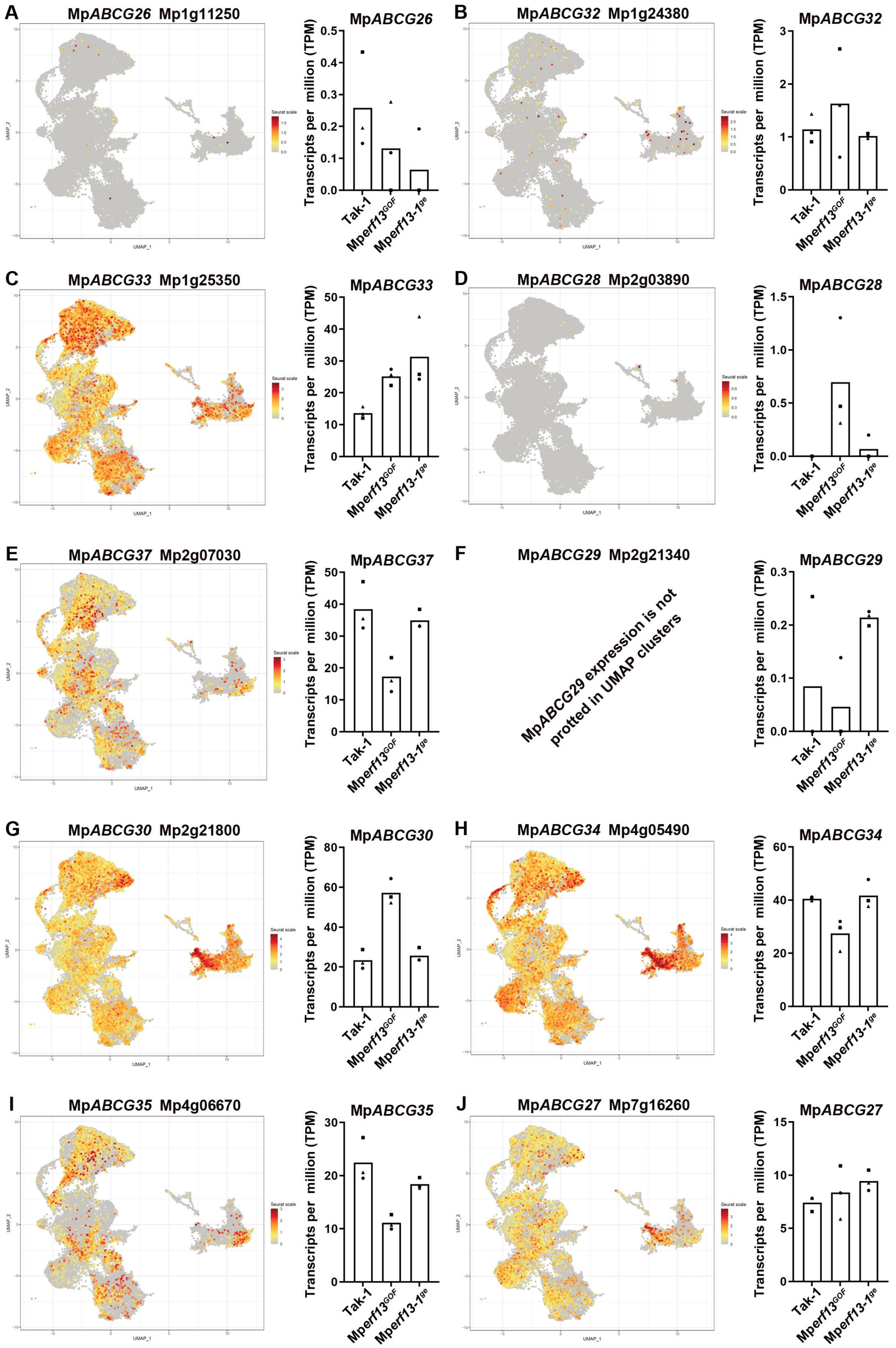


**Supplementary Figure S2. Transcriptional expression profiles of all *ABCG* genes in *M. polymorpha*.** Expression levels of all *ABCG* genes are shown as UMAP cluster distributions generated using Single Cell Browser (Wang et al., 2023) and transcripts per million (TPM) values re-calculated from the RNA-sequencing data of Kanazawa et al. (2020). Data for Mp*ABCG1* and Mp*ABCG36* are presented in Figure 3 and Supplementary Figure S1, respectively. Full and half *ABCG* genes are presented in panels (A–K) and (L–AH), respectively.

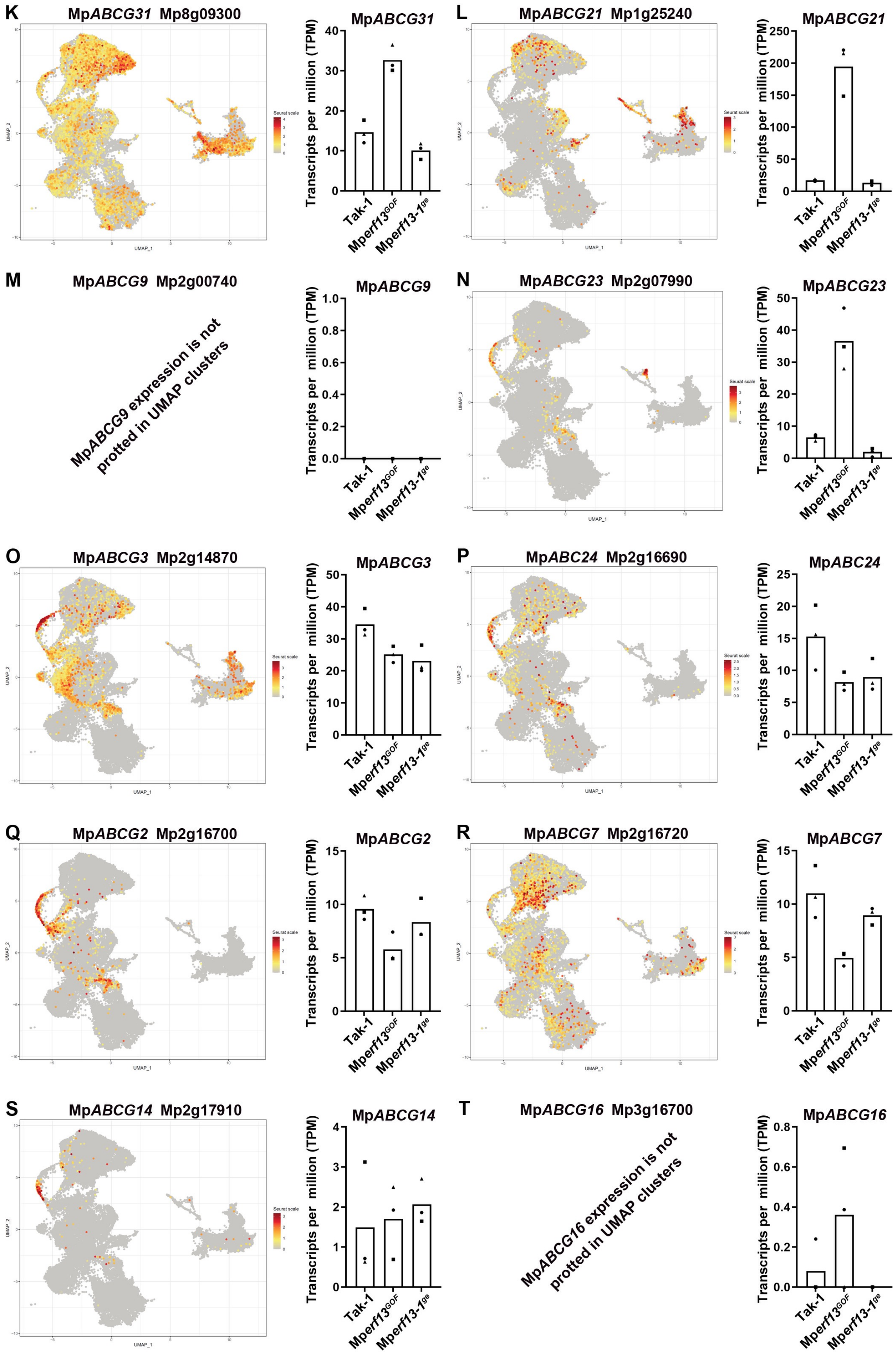


**Supplementary Figure S2. (Continued).**

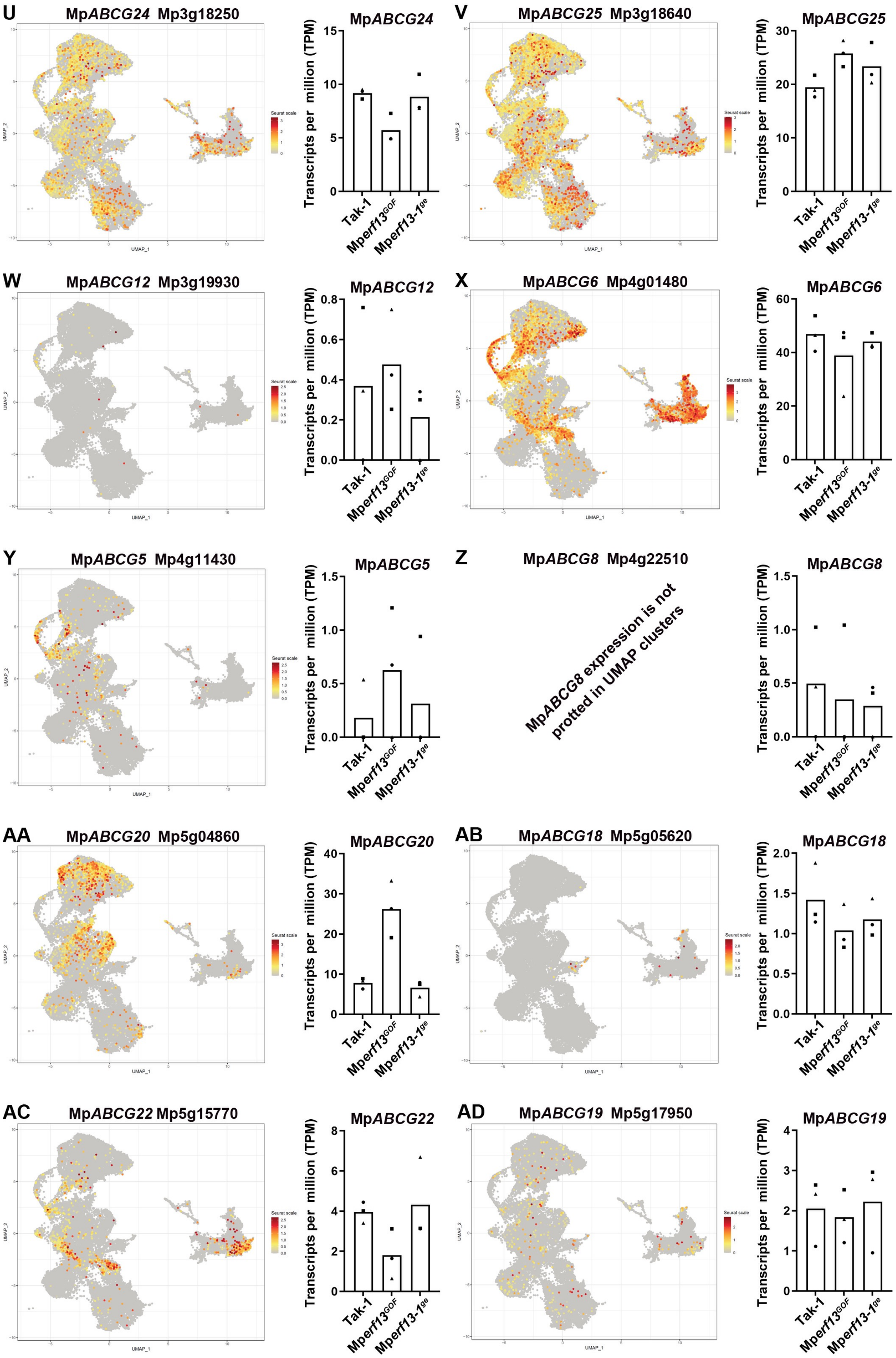


**Supplementary Figure S2. (Continued).**

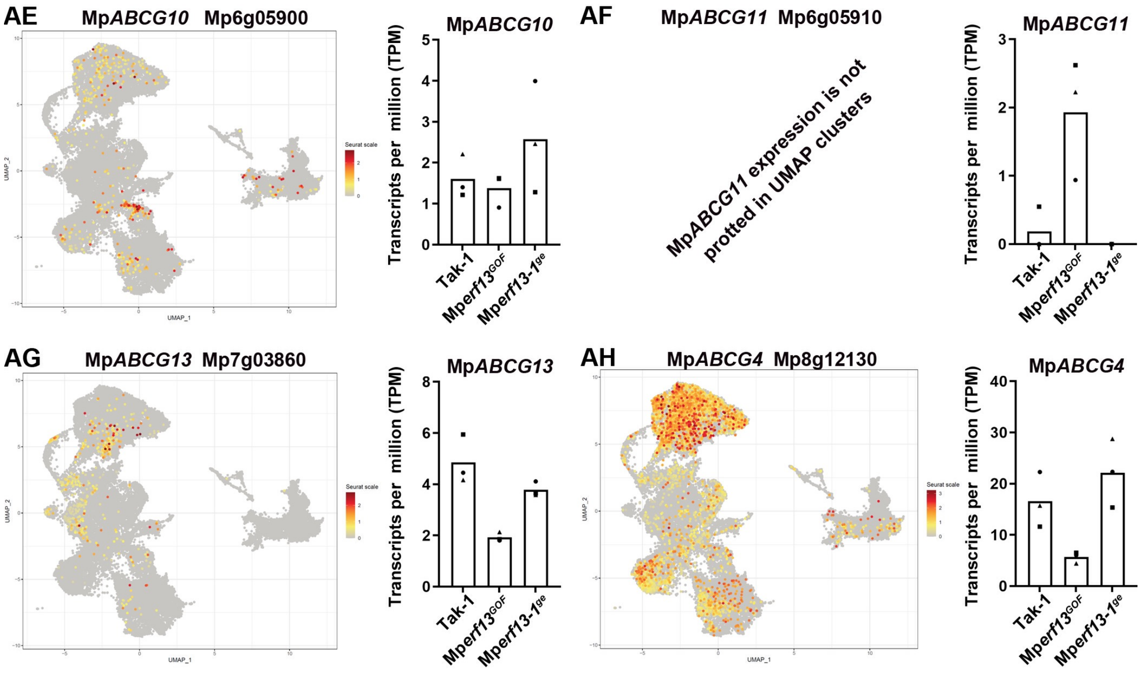


**Supplementary Figure S2. (Continued).**

### 1. D-Limonene (94)

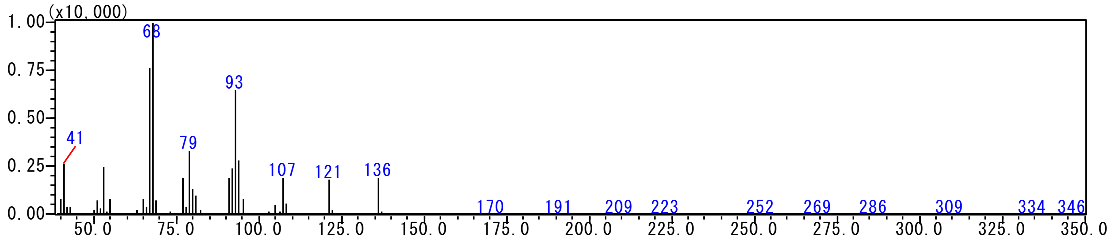


### 2. 1-Octen-3-yl-acetate (94)

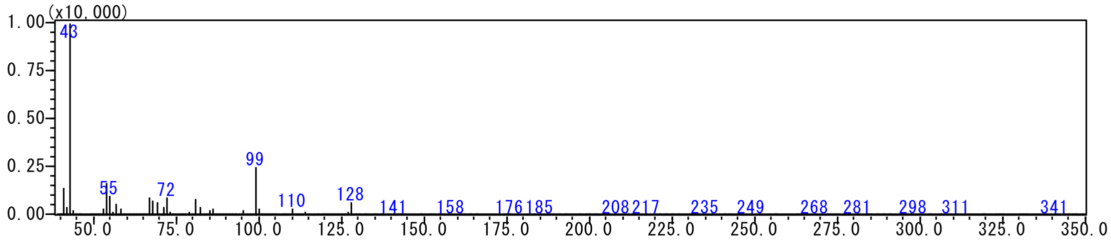


### 3. α-Chamigrene (89)

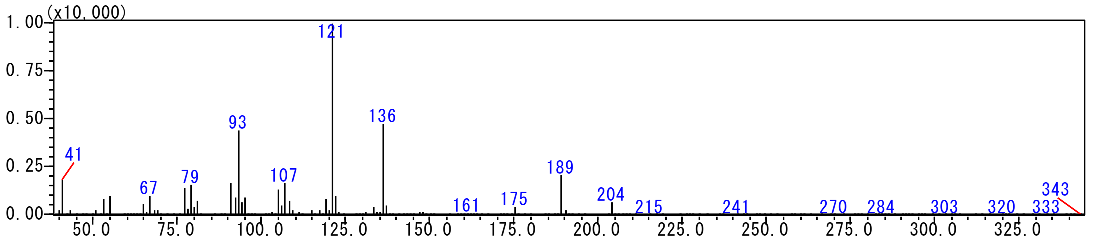


### 4. Thujopsene (96)

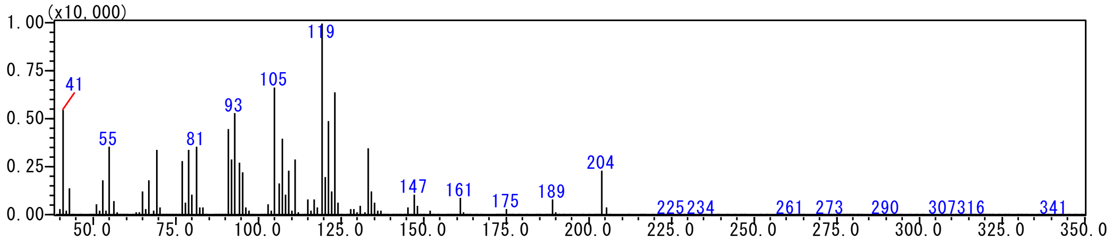


**Supplementary Figure S3. Mass spectra of compounds 1–9.**
For each compound, the most similar match in the National Institute of Standards and Technology (NIST)-17 library is shown, with the similarity score indicated in parentheses.

5. β-Chamigrene (95)

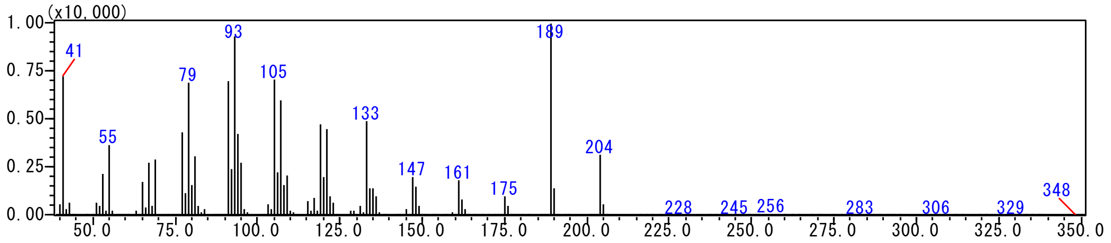


6. *cis*-α-Bisabolen (92)

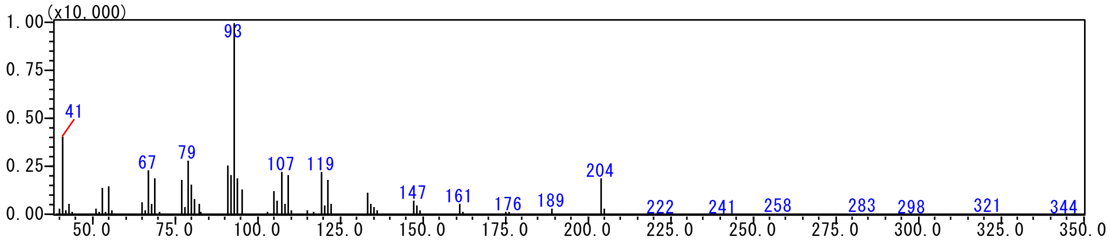


7. β-Himachalene (95)

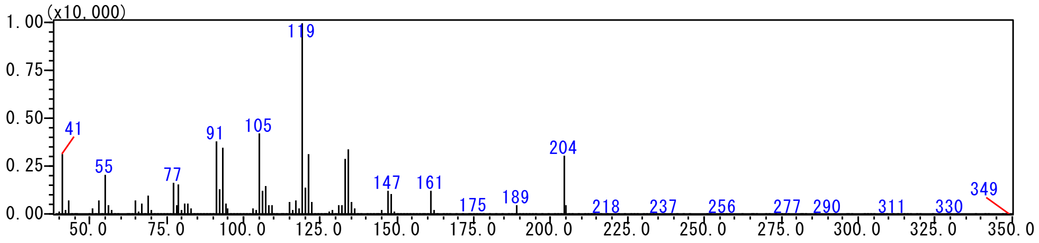


8. 1,1,3-Trimethyl-3-(2-methyl-2-propenyl)cyclopentane (84)

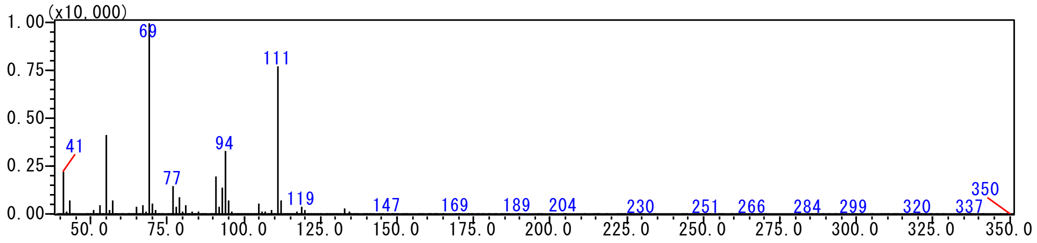


9. Caryophyllene oxide (87)

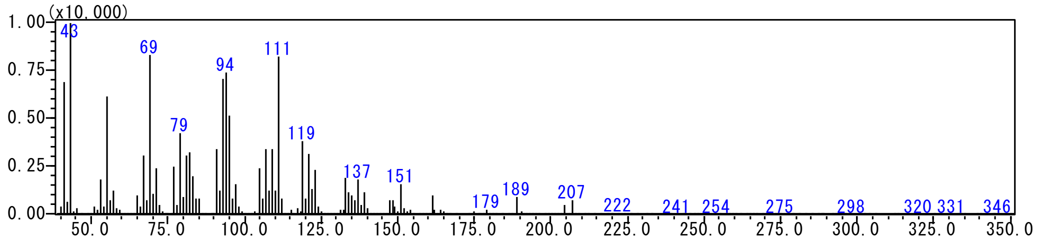


**Supplementary Figure S3. (Continued).**

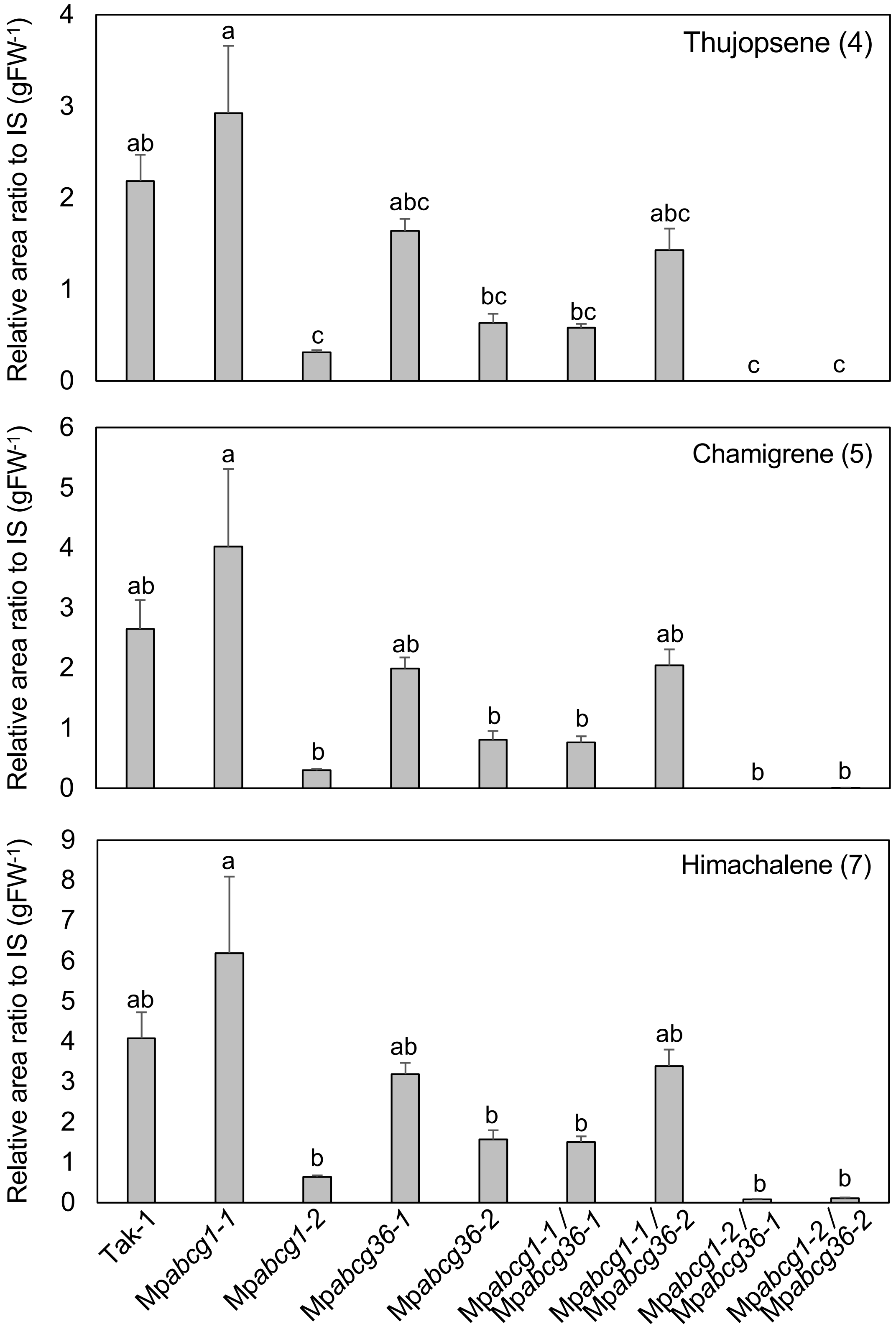


**Supplementary Figure S4. Levels of sesquiterpenoids in all mutants generated in this study.**
Levels of three sesquiterpenoids identified in the extracts of the six-week-old thalli of all mutants generated in this study are shown. The relative peak area of each compound normalized to the internal standard (IS, tetralin) is shown. Data are represented the mean ± SE of three biological replicates. Different lowercase letters indicate the significant differences (one-way analysis of variance [ANOVA], followed by Tukey's test; $P < 0.05$).

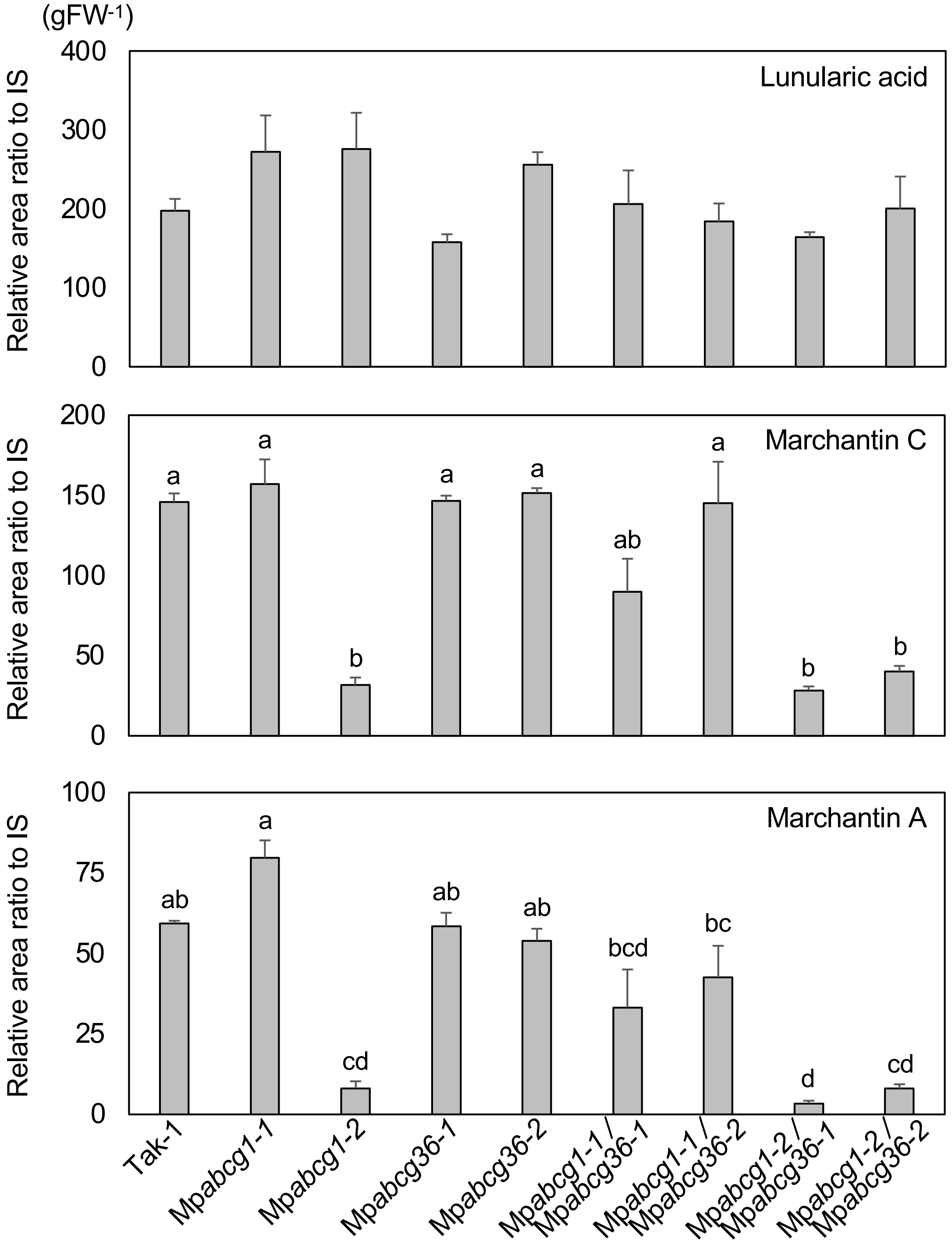


**Supplementary Figure S5. Levels of three bibenzyls in all mutants generated in this study.**

Levels of three bibenzyls identified in the extracts of the six-week-old thalli of all mutants generated in this study are shown. The relative peak area of each compound normalized to IS (nordihydroguaiaretic acid [NDGA]) is shown. Data are represented as the mean ± SE of three biological replicates. Different lowercase letters indicate the significant differences (one-way ANOVA, followed by Tukey's test; $P < 0.05$).
